# Phylogenomic analysis of the collagen-like BclA proteins in *Clostridioides difficile*

**DOI:** 10.1101/2025.06.24.661383

**Authors:** Francisca Cid-Rojas, Enzo Guerrero-Araya, Christian Brito-Silva, Marjorie Pizarro-Guajardo, César Rodríguez-Sanchéz, Daniel Paredes-Sabja

## Abstract

*Clostridioides difficile* is a Gram-positive, anaerobic, spore-forming bacterium and a major nosocomial pathogen, notable for its high genetic diversity. *C. difficile* has been classified into five classical phylogenetic clades (C1 to C5) and five cryptic clades (C-I to C-V), reflecting its extensive genome plasticity. In addition, *C. difficile* spores are considered essential for the onset, persistence and transmission of the disease, and their exosporium layer has hair-like projections formed by the BclA-family of proteins (BclA1, BclA2 and BclA3). Previous work in C1 and C2 strains has demonstrated that the absence of these proteins affects spore germination, pathogenesis, persistence, and recurrence of CDI. Nevertheless, the conservation of BclAs across different *C. difficile* clades remains unclear. In this work, genomics analysis in more than 25,000 *C. difficile* genomes revealed that the prevalence and variability of the BclAs was not conserved across classical and cryptic clades. The most represented clade on the dataset was C1, where roughly 50% of the genomes possessed all three *bclA* genes. Pseudogenization of *bclA1* or all *bclAs* was observed in C2 and C3, respectively. Additionally, the absence of *bclA1* in C4 and of both *bclA1* & *bclA3* in C5, further demonstrates the divergence of BclAs among *C. difficile* clades. Subsequent analysis revealed high variability in the central collagen-like region (CLR) of all three BclAs, and a highly conserved *bclA1* pseudogenization event in most members of C2. Overall, the extensive variability of *bclA*, attributed to the CLR, prevalence of a *bclA1* pseudogenization and complete absence of *bclAs* in certain clades, is likely to impact spore morphogenesis and pathogenesis across different *C. difficile* clades.

## Introduction

*C. difficile*, a ubiquitous anaerobic spore-producing microbe, is frequently implicated in nosocomial or antimicrobial-associated infections, collectively known as *C. difficile* infections (CDI) ^1^. This environmentally persistent pathogen is capable of causing a wide range of symptoms, from mild diarrhea to severe pseudomembranous colitis ^2^. As a leading cause of healthcare-associated infections, *C. difficile* poses a significant burden on global health, with incidence rates ranging from 2 to 8 cases per 10,000 patient-days ^3, 4^. Conventional treatment approaches for CDI involve antibiotics, such as vancomycin, metronidazole, and fidaxomicin ^5^. However, despite these interventions, approximately 15 to 35% of CDI cases experience recurrence (R-CDI) ^6^.

*C. difficile*’s infection lifecycle begins with the germination of endogenous or recently acquired C. *difficile* spores ^7^. Two main virulence factors, the toxins TcdA and TcdB, are primarily responsible for the clinical manifestations of *Clostridioides difficile* infection (CDI) ^8, 9^. These toxins monoglucosylate host Rho family GTPases, leading to downstream effects such as tight junction disruption, epithelial cell detachment, and compromised intestinal barrier function^8, 9^. Additionally, toxin exposure prompts intestinal epithelial cells to secrete pro-inflammatory cytokines, which attract neutrophils and initiate acute mucosal inflammation^8, 9^. This inflammatory response further activates immune cells, intensifying tissue damage.

*C. difficile* spores are the main virulence factor related with disease recurrence ^10, 11^. During CDI, *C. difficile* begins a sporulation cycle leading to the formation of newly dormant spores^10^, that interact with host surfaces^12–15^, and persist to cause R-CDI^16^. Essential in these interactions are the hair-like projections on the spore surface, where the collagen-like BclA3 protein is essential for their formation^16^. Two additional BclA orthologues (BclA1 and BclA2) are encoded in some *C. difficile* strains. Work in strains 630 (clade 1) and R20291 (clade 2), shows their implication in spore-host interactions, colonization and recurrence of CDI ^16, 17^.

*C. difficile* has tremendous genetic diversity ^18, 19^, where its core genome represents only 10-20 % of its pan-genome. This likely contributes to *C. difficile* adaptation to a wide host-range and persist in environmental reservoirs^20^. This diversity is evidenced in the five distinct phylogenetic classical clades and five cryptic clades, based on multi-locus sequencing and core-genome phylogeny^18^. While Clade 1 is the most represented clade, with the broadest geographic distribution ^3, 21^, clade 2 includes epidemically relevant strains of ribotype 027 (RT027) lineage and mainly associated with hospital outbreaks ^22–24^. Clade 3 are uncommon yet phenotypically share similarities with Clade 2 strains^25^. Clade 4 includes strains that are often clindamycin- and fluoroquinolone-resistant, and associated with outbreaks in Asia, North America and Europe^26^. Finally, Clade 5 is the most genetically distant clade from all classical clades and based on its 96% average nucleotide identity (ANI) to all other four clades, it is hypothesized to be an early predecessor of all four clades^27^. However, several genomospecies, with ANI values lower than 95% to all 5 classical clades and with highly divergent toxin gene architecture have been identified and defined as cryptic clades (CI to V)^3, 18, 28, 29^.

The outermost layer of the spores, called exosporium, possess surface-proteins resembling hair-like projections^11, 30^. Transmission electron microscopy (TEM) of the spores has revealed that exosporium morphology seems to be strain-dependent, as spores from the epidemically relevant strains (including R20291) have a hair-like nap, meanwhile 630, a strain widely used in scientific research, has a compact exosporium layer without hair like structures^11^. These hairs resemble those present on spores from members of the *B. cereus* group (*B. anthracis*), which have been determined to be formed by BclA glycoproteins^31, 32^. Strong *et al.*^33^ identified three *bclA* homologues on the genome of *C. difficile*; *bclA1, bclA2* & *bclA3*

Bacterial proteins like *B. anthracis* BclA & BclB and *Streptococcus pyogenes* (*S. pyogenes*) Scl1 & Scl2, have been thoroughly characterized previously^31, 32, 34–39^. These proteins were encountered to be displayed on the outermost layer of their respective bacterial morphotype with a central triple-helix rod shape structure composed of canonical GXY collagen repeats^40^ and a globular end terminal domain^32, 36, 38, 39^, thus they are referred to as collagen-like proteins. Although they differ in length, the domain distribution, N-terminal domain (NTD), collagen-like region (CLR), and C-terminal domain (CTD), is the same as *C. difficile* exosporium BclA proteins (**Fig. 1A**).

**Fig. 1.**
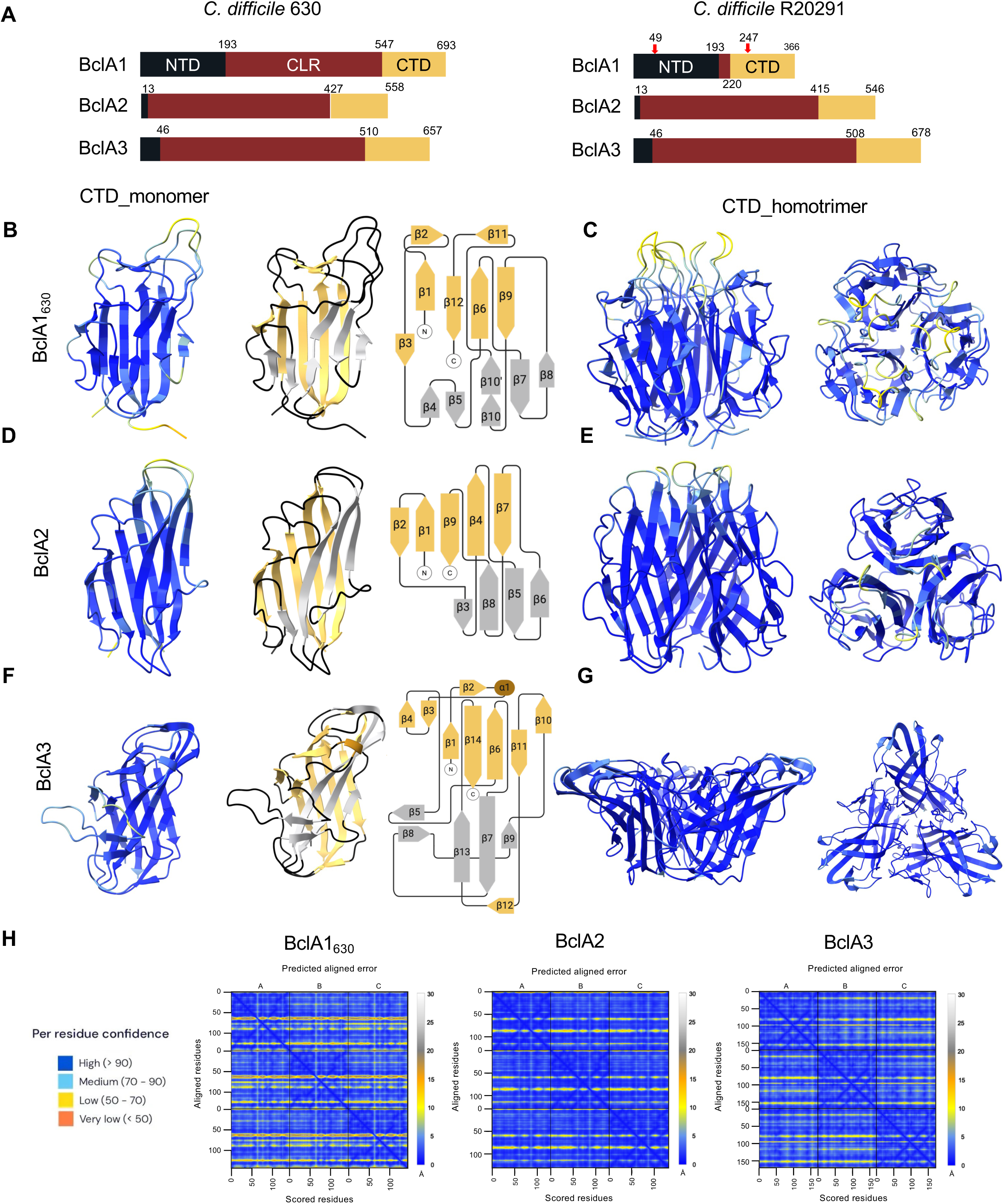
BclA domain structure in *C. difficile* 630 & R20291 strain and AlphaFold3 BclA CTD predictions. **(A)** Schematic representation of *C. difficile* 630 and R20291 BclA1, BclA2 and BclA3 protein sequences. NTD (Black): N-terminal domain, CLR (Maroon): Collagen-like region, CTD (Yellow): C-terminal domain. In red arrows are depicted two stop codons that led to early termination of translation of BclA1_R20291_. **(B,D,F)** AlphaFold3 prediction of CTDs monomers of BclA1_630_, BclA2_R20291_ and BclA3_R20291_, and topology models to the right are shown. *α*-helix and loops are displayed in brown and black, respectively. Yellow *β*-sheets are located on the inner face of homotrimers models. Grey *β*-strands are located on the outer face of homotrimers models. Encircled “N” and “C” represent the amino- and carboxyl-terminus, respectively. Structural predictions indicate that BclA1_630_ had a total of 12 *β*-sheets, BclA2_R20291_ had a total of 9 *β*-sheets and BclA3_R20291_ encodes 14 *β*-sheets and 1 *α*-helix. **(C,E,G)** AlphaFold3 prediction of CTDs homotrimers of BclA1_630_, BclA2_R20291_ and BclA3_R20291_ and **(H)** their Predicted aligned error (PAE) are shown.

Despite the wealth of genomic data for *C. difficile*, there is a lack of knowledge on the diversity of spore surface proteins implicated in pathogenesis. Therefore, in this work we explored BclA prevalence and variability in more than 25,000 *C. difficile* genomes and used to generate a core-genome phylogenetic tree, where four new clades were introduced, representing the complex and changing taxonomical organization of this pathogen; the tree was divided into classical clades (C1 to C7) and cryptic clades (C-I to C-VII). Subsequent *bclA* locus search and analysis revealed a high variability of the central collagen-like region (CLR) of the proteins, indistinctively of the clades to which they belong, and the finding of a conserved *bclA1* pseudogenization event present across most members of C2, which was not observed for *bclA2* or *bclA3.* Complementation with a full length *bclA1* from 630 strain (C1) to R20291 strain, which is a member of C2, resulted in an increase of exosporium and hair-like projection’s length. Overall, the extensive variability of *bclA*, mainly attributed to their CLR, prevalence of a *bclA1* pseudogenization and even the complete absence of *bclAs* in certain clades.

## Materials and Methods

### Genome collection

The complete dataset from Enterobase was obtained, which included 25,145 carefully selected *C. difficile* genomes, as well as diverse metadata, with a cutoff date of September 30, 2022. This dataset is a valuable resource that encompasses information on a wide variety of strains and conditions. In addition to the data provided by Enterobase, 21 additional genomes were incorporated from suspected cryptic isolates, with the aim of further expanding the diversity and scope of the analysis.

### Genomes assemble

Genomes were assembled, annotated, and evaluated using a pipeline ^41^ comprising TrimGalore v0.6.5, SPAdes v3.6.043, Prokka v1.14.5, and QUAST v2.344. Next, Kraken2 v2.0.8-beta was used to screen for contamination and assign taxonomic labels to reads and draft assemblies.

### Multi-locus sequence typing

The assembled genomes underwent a thorough analysis for Multi-locus sequence typing (MLST) utilizing FastMLST v0.0.15 ^42^ to identify any significant genetic variations. During this process, new alleles and sequence types (STs) were detected and subsequently verified by submitting the assembled contigs to the PubMLST database (https://pubmlst.org/cdifficile/) ^43^, a widely recognized public resource for molecular typing and microbial genome diversity research

### Core-genome genes and recombination-free phylogeny

To identify core-genome genes in the analysis, DIAMOND blastp ^44^ search was employed, coupled with a bidirectional best hit approach, using the proteins predicted by Prodigal ^45^ from both isolated and predicted proteomes of *C. difficile* 630 as query sequences. A gene was designated as a core gene if it was present in at least 99% of the isolates. Following the identification of core genes, MAFFT ^46^ was used to align each gene individually, and subsequently concatenate the alignments to form a comprehensive dataset.

To account for the potential influence of recombination events on the analysis, it was applied ClonalFrameML ^47^ to remove any signs of recombination from the concatenated alignment. With a recombination-free alignment at hand, a phylogenetic tree was constructed using IQ-tree ^48^, implementing 1000 ultrafast bootstrap replicates to assess the robustness of the inferred tree topology.

Upon completion of the phylogenetic tree, final clade delimitations were stablished, which allowed to group *C. difficile* isolates into distinct evolutionary lineages. This comprehensive approach to genome analysis and phylogenetic tree construction provides valuable insights into the genetic relationships and evolutionary history of *C. difficile* isolates in the dataset used.

### 3D structural inference

Alphafold3^49^ was employed, a state-of-the-art protein folding prediction algorithm, to determine the three-dimensional structure of the proteins under investigation. The Alphafold3 system utilizes machine learning techniques to predict protein structures with remarkable accuracy, providing valuable insights into protein function and molecular interactions.

Predictions were performed with 24 iterative cycles (seeds). Running multiple seeds of the model helps to refine the prediction at each cycle, enhancing the overall reliability and accuracy of the predicted structures. The highest-ranking predicted structure (rank 0), that had the lowest predicted alignment error (PAE) was selected for further analysis. The predicted alignment error provides a quantitative measure of the accuracy of the predicted structure compared to the native structure.

### Sequence alignment

Clustal Omega ^50^, a widely recognized and efficient tool for generating multiple sequence alignments, was used to align the sequences of interest. Clustal Omega is an advanced algorithm that excels in aligning both nucleotide and amino acid sequences, making it a popular choice for comparing homologous sequences and identifying conserved regions, which can provide crucial insights into protein function, structure, and evolutionary relationships. To perform the multiple sequence alignments, default parameters provided by the Clustal Omega software were used.

EMBOSS Needle, was used to perform pairwise sequence alignment. It creates an optimal global alignment using the Needleman-Wunsch algorithm. Default parameters provided by the software were used.

### Extraction and distribution of BclAs proteins

To extract the sequences of the three BclAs of interest, each genome in the dataset was first annotated using Prodigal ^45^, a widely used and accurate gene prediction tool designed for microbial genomes. This allowed to identify the coding sequences within each genome and obtain a comprehensive set of gene annotations for further analysis. Following the genome annotation step, a search for the upstream and downstream genes of each BclA within the annotated genomes using DIAMOND ^44^ was performed. By locating these flanking genes, it was possible to define the genomic region encompassing each BclA. Subsequently, these regions were extracted for further investigation and analysis. To ensure accuracy and precision in the extraction process, this final step was performed manually. Given the large number of isolates in the dataset, the focus was set on single nucleotide polymorphisms (SNPs) present in the alignment for constructing the core-genome phylogenetic tree. To achieve this, the program employed was RapidNJ (https://github.com/somme89/rapidNJ), a fast and efficient algorithm for constructing phylogenetic trees based on the Neighbor-Joining method. To minimize redundancy in the dataset and improve computational efficiency, deduplication using CD-HIT ^51^ was performed, a widely utilized clustering tool that identifies and removes duplicated sequences based on sequence similarity thresholds. This step ensured that the dataset contained only unique sequences, enabling more accurate and meaningful downstream analyses. Lastly, custom Python scripts were used to generate schematic representations of the results. These visualizations provided a clear and concise representation of the findings, facilitating the interpretation and communication of the insights gleaned from the study.

## Results

### Structural modeling of *C. difficile* BclA proteins

*C. difficile* encodes at least three orthologues of the BclA family of collagen-like proteins^52^. *C. difficile* strains 630 (Ribotype 012, clade 1) and R20291 (Ribotype 027, clade 2), despite belonging to different clades, they possess similar BclA proteins and share almost identical genetic context. Amino acid sequence comparison reveals that BclA2 of 630 and R20291 have ∼ 95 % identity (**Fig. S3, S4**), while BclA3 shares ∼ 87 % identity (**Fig. S5, S6**). However, in the case of BclA1, identity dropped to ∼ 51 %, however reaches 97 % upon comparing only the NTDs (**Table S1**). This is due to the pseudogenization of *bclA1* in R20291 strain by two nonsense mutations at: i) nucleotide positions A145T, resulting in an early-stop codons that lead to formation of a small 48 amino acid NTD polypeptide; and ii) at nucleotide position C739T which yields an additional stop codon (**Fig. 1A, S1, S2**). When the alignment was performed by individual BclA domains, gap-free alignment regions revealed more than 93% identity was observed for NTD and CTDs of BclA2 and BclA3, except for the CLR of BclA3 with 77% identity (**Table S1**). As to better compare both BclA1 from 630 and R20291 strain from a protein level in further analysis, and since *bclA1_R20291_* is pseudogenized but the downstream DNA sequence follows the same structure as a canonical *bclA1* (**Fig. S1**), the two nonsense mutations were repaired to obtain a full length repaired BclA1_R20291_ (rBclA1_R20291_). Overall, BclAs of both strains were alike with the CLR domain having the least similarity.

**Fig. 2.**
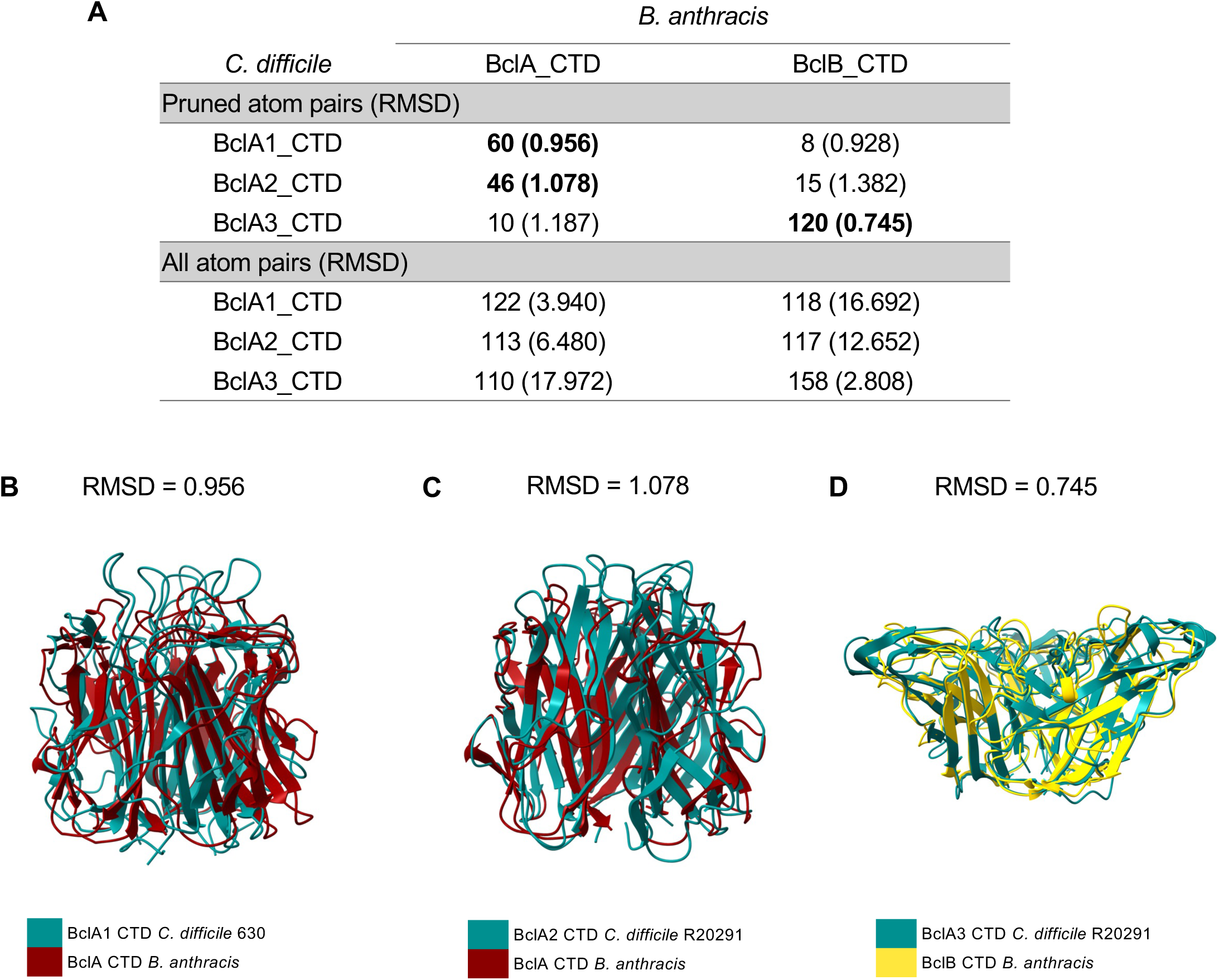
*C. difficile* BclA comparison with BclA and BclB CTD domain from *B. anthracis*. Alphafold3 predicted CTD homotrimers structures of *C. difficile* (Cd) and *B. anthracis* (Ba) were superimposed on ChimeraX (v.1.9) using “matchmaker” command with default parameters. **(A)** RMSD for both pruned and all atom pairs is reported for all combinations possible. Numbers in bold represent the lowest RMSD achieved for each protein comparison. Superimposed pairs are **(B)** Cd_BclA1_CTD_630_ with Ba_BclA_CTD, **(C)** Cd_BclA2_CTD_R20291_ with Ba_BclA_CTD and **(D)** Cd_BclA3_CTD_R20291_ with Ba_BclB_CTD. *C. difficile* 630 genome used as reference was GenBank ID: AM180355.1. *C. difficile* R20291 genome used as reference was GenBank ID: CP029423. *B. anthracis* genome used as reference was GenBank ID: AE016879.1. RMSD: Root mean square deviation, Cd: *Clostridioides difficile*, Ba: *Bacillus anthracis*.

Given the differences in BclA1 and similarities in BclA2 and BclA3 between strains 630 and R20291, AlphaFold3 (AF3) ^53^ was used to create structural predictions of full-lengths (fl) BclA1_630_, rBclA1_R20291_, BclA2_R20291_ and BclA3_R20291_. Initial monomeric predictions for all fl-BclAs show low confidence models for almost the entire protein. Upon modeling the individual domains, monomeric CLR of all four proteins proved to be disorganized with very low predicted confidence. However, monomeric CTDs of BclA2 and BclA3, except for BclA1_630_ and rBclA1_R20291_, were confidently predicted as globular structures with low expected positional error. Interestingly, the NTDs of BclA1_630_ and rBclA1_R20291_ were predicted to be globular domains, while the NTDs of BclA2 and BclA3 had not predicted secondary structure (**Fig. S7A, S7B - S10A)**. Among these NTDs, rBclA1_R20291_ was predicted with higher confidence than that of BclA1_630_, likely due to the longer CLR domain in BclA1_630_ that could be affecting AF3 predictions (**Fig. S7A & S7B**). Overall, monomeric predictions of all BclA were confident mainly at the CTD, while failing to accurately predict NTD and CLR.

Collagen proteins have a globular trimerization domain at one of their terminal domains which together with their intrinsic GXY repeats, are driving forces for the formation of triple-helix structures^49, 54^, suggesting that all three BclA are likely to form stable predictable trimers. Strikingly, although homotrimers were predicted for all fl-BclAs, the confidence of these interactions was limited mainly to the CTDs, where BclA2 had the highest predicted homotrimerization confidence, forming a tight CTD-protein complex (**Fig. S8B**), followed by BclA3 (**Fig. S9B**). By contrast, both BclA1_630_ and rBclA1_R20291_ had an apparent low confidence homotrimerization prediction (**Fig. S7C & S7D**). Unfortunately, AF3 was unable to predict the triple-helix structure in all *C. difficile* the collagen-like BclA proteins, likely due to limitations of the model.

The shortest collagen-like proteins that have structural properties of collagens are synthetic collagen mimetic peptides (CMPs), which have at least 6 GXY repeats that can self-assemble into triple-helix structures^55, 56^. Therefore, a truncated CLR with 10 GXY repeats was created in all three BclA to evaluate their predictability using AF3. The predicted structures of 10 GXY BclAs improved compared to full-length BclAs. Prediction of 10 GXY versions of BclA2 and BclA3 revealed a more accurate organization of BclA trimers, comprising of an N-terminal domain, a truncated central collagen-like region, and a C-terminal domain (**Fig. S10**). However, for trimerization of BclA1_630_, although prediction of truncated BclA1 improved compared to full length, trimerization of the CTD was not as confident as for truncated versions of BclA2 and BclA3 (**Fig. S10**). Altogether, structural predictions strongly support that the CTDs of BclA2 and BclA3 tend to interact and are likely trimerization domains. By contrast, BclA1 trimerization had no confident prediction, even in the truncated BclA1 versions.

Collagen molecules bind to different types of collagens forming heterotrimeric proteins^49, 57, 58^. Therefore, to test the hypothesis that *C. difficile* BclA orthologues could form heterotrimers, AF3 predictions were employed to predict the structure of trimeric complexes composed of different BclA proteins. Combination of distinct BclA sequences were input as separate chains, allowing AF3 to model potential inter-chain interactions and assess compatibility in forming stable trimeric assemblies. Resulting predictions revealed no expected interactions between the CTD of different BclA, even though most molecules were predicted with high confidence as independent units: i) CTD_BclA2-BclA3_R20291_ (**Fig. S11**), and ii) CTD_BclA1-BclA2-BclA3_630_ (**Fig. S12**). These results support the notion that *C. difficile* BclAs do not form heterotrimeric protein complexes.

Further analysis of CTDs, demonstrates that BclA1_630_, BclA2 and BclA3 is formed by total of 12, 9 and 14 predicted beta sheets, while only CTD of BclA3 has an alpha helix in between beta-sheets 2 and 3 (**Fig. 1B, 1D, 1F**). While they all form a homotrimer barrel-type structure predicted with high confidence, BclA1_630_ and BclA2 have a ball-like shape, and BclA3 a trapezoid-like shape that, when viewed from above, forms an isosceles triangle. Interestingly, the areas of lowest confidence in the inference of the three-dimensional structure are in the coils at the tip of each monomer conforming the structures (**Fig. 1C, 1E, 1G, 1H**). These structural differences suggest that BclA1, BclA2, and BclA3 are homologues of each other and may have different functions in the exosporium.

Structural similarities between *C. difficile* and *B. anthracis* BclA proteins.

To investigate the relationship between *C. difficile’s* BclAs and *B. anthracis’s* collagen-like proteins (BclA and BclB), AF3 was used to model the protein’s CTD_homotrimers of both bacteria. *C. difficile* BclA_CTD domains were independently superimposed to *B. anthracis* BclA_CTD and BclB_CTD using ChimeraX (**Fig. 2A**). Calculation of RMSD (Root Mean Square Deviation) was used to quantify atomic differences between the superimposed proteins. Both, BclA1_CTD_630_ and BclA2_CTD_R20291_ had the best hit with BclA_CTD of *B. anthracis*, with an RMSD of 0.956 and 1.078, respectively (**Fig. 2B and 2C**). On the other hand, BclA3_CTD_R20291_ showed a high structural similarity with BclB_CTD *B. anthracis*, with an RMSD of 0.745 (**Fig. 2D**). Additionally, low sequence identity was observed between these domain (15.8-19.5%) and a similarity between 23.9-35.6% (**Fig. S13**). These results suggest that despite the low identity between the CTDs of *C. difficile* and *B. anthracis*, they share comparable predicted structures.

### Phylogenomic distribution of BclA across the *C. difficile* lineage

To further explore the diversity of BclA among *C. difficile* lineage, a set of 25,165 *C. difficile* genomes were analyzed for the presence of these proteins (**Table S2**). BclA1 was found in 15,680 (**Table S3**), BclA2 in 21,540 (**Table S4**) and BclA3 in 17,685 genomes (**Table S5**). The distribution of hits in a phylogenetic core-genome tree is shown in Figure 3A. In the case of BclA1 occurrence, a homogeneous distribution is observed in all clades except for clade 4 and clade 5, where BclA1 was not found in most or any of the genomes searched, 74% and 100%, respectively (**Fig. 3A, S14A, S14B & Table S6**). The presence of BclA2 was homogeneous across all clades, the same as BclA3 except for clade 5, where it was not found in 97.6% of the genomes (**Fig. 3A, S14B & Table S6**).

**Fig 3.**
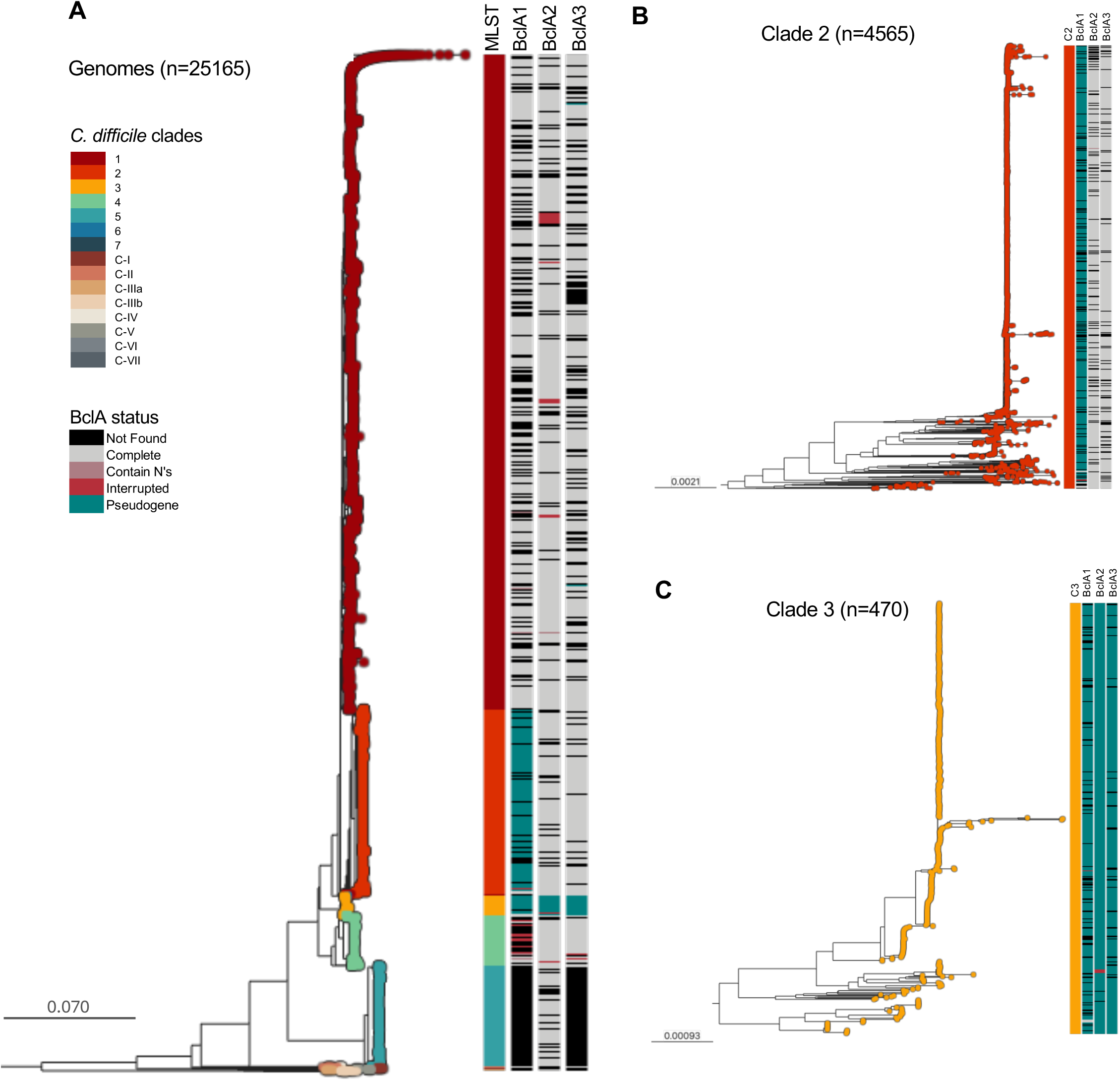
Distribution of BclAs in known diversity of *C. difficile*. **(A)** Phylogenetic tree of 25k core-genome of *C. difficile*, showing the clade each genome belongs to. Horizontal bars represent the state of BclA in a particular genome (not found, complete, containing Ns, interrupted, or pseudogenized). **(B)** Zoomed-in view of the Clade 2 subtree. **(C)** Zoomed-in view of the Clade 3 subtree

Clade 1 comprised 16,264 genomes, among which the majority harbored complete open reading frames (ORFs) for *bclA* orthologues: *bclA1* in 68.42%, *bclA2* in 85.75%, and *bclA3* in 74.02% of genomes. Notably, 54.65% (n = 8,889) of genomes in this clade contained all three *bclA* homologs. A smaller subset presented only *bclA2* (10.25%), *bclA3* (3.24%), or *bclA1* (1.53%) individually. Pseudogenized *bclA* genes were rare in this clade, with *bclA1*, *bclA2*, and *bclA3* pseudogenization observed in only 0.1%, 0.2%, and 0.17% of genomes, respectively. These results indicate that Clade 1 harbors a largely intact and widespread distribution of *bclA*, with minimal gene inactivation (**Fig 3A, Table S6-S8**).

Of the 4,565 genomes analyzed of Clade 2, only 56 genomes (1.23%) contained a complete *bclA1* ORF. In contrast, 80.92% (n = 3,694) exhibited a pseudogenized *bclA1*, indicating a clade-wide inactivation of this gene. Despite this, most genomes (69.31%) retained all three *bclA* homologs, including pseudogenized copies. A smaller fraction harbored only one *bclA* ORF (*bclA1*, 2.85%; *bclA2*, 3.5%; *bclA3*, 2.8%), and pseudogenized *bclA2* and bclA3 were found in 0.18% and 0.11% of genomes, respectively. These findings suggest that while *bclA1* is frequently inactivated in Clade 2, the overall *bclA* repertoire remains largely preserved (**Fig 3B, Table S6-S8**).

Among the 470 genomes analyzed from Clade 3, pseudogenization was widespread, with 81.7% of genomes containing at least one inactivated *bclA* gene. Complete ORFs for *bclA2* and *bclA3* were absent from all genomes, while only three genomes carried a complete *bclA1* (0.6%). Pseudogenized alleles were prevalent: *bclA1* in 84.04%, *bclA2* in 98.72%, and *bclA3* in 93.83% of genomes. Only a small subset contained a single *bclA* gene (*bclA2*, 4.26%, n=20; *bclA3*, 0.21%, n=1), and none harbored *bclA1* alone. This suggest that Clade 3 might not have functional BclA proteins, based on its highly pseudogenized alleles (**Fig 3C, Table S6-S8**).

Clade 4 included 1,254 genomes. A majority possessed complete ORFs for *bclA2* (90.59%) and *bclA3* (82.62%), while *bclA1* was found intact in only 25.52% of genomes. However, just 17.15% (n = 215) of genomes contained all three *bclA* full length homologs. A minority carried only *bclA1* (0.88%), *bclA2* (6.94%), or *bclA3* (4.55%), and pseudogenization was limited, occurring in 0.32% (*bclA1*), 0% (*bclA2*), and 0.16% (*bclA3*) of genomes. These data show that while *bclA2* and *bclA3* are well-conserved in Clade 4, *bclA1* is frequently absent or incomplete, limiting the presence of the full paralog set (**Fig S14A, Table S6-S8**).

Out of the 2,491 genomes of Clade 5, 79.69% possessed a complete *bclA2* ORF. Interestingly, the majority was missing *bclA1* (100%) or bclA3 (97.59%), and a few genomes *bclA2* (19.55%). Pseudogenized alleles were scarce and included *bclA2* in 0.36% and *bclA3* in 0.04%. Overall, these results suggest that Clade 5, with the complete absence of *bclA1* and *bclA3*, will only have BclA2 protein in its repertoire. (**Fig S14B, Table S6-S8**).

Unfortunately, the *C. difficile* data set had few representatives of Clade 6 (n = 5) and Clade 7 (n = 1), therefore it is not possible to determine trends or statistics for these clades (**Table S6**). Regarding the cryptic clades, almost the entirety of C-IIIa and C-IIIb genomes where missing *bclA1* and *bclA3*, and roughly 50% did not presented *bclA2*. Meanwhile, C-II and C-IV genomes did not present *bclA2* at all, contrary to C-V where 43% only presented *bclA2* (**Fig. S15C, Table S6, S7, S9**).

Pseudogenization of *bclAs* across *C. difficile* clades.

As stated before, pseudogenization of *bclA1* is present in 80.92% of Clade 2 and 84% of Clade 3 genomes (**Fig. 4A, Table S6-S8**). In the case of *bclA2*, 98.72%, and for *bclA3*, 93.83% of Clade 3 genomes have these genes pseudogenized (**Fig. 4B, 4C & Table S6-S8**). Notably, specific pseudogenized events were highly abundant for each of the *bclAs*. At the sequence level, *bclA1* pseudogenization occurred at residue 49 in 99.49% of cases, consistent with the pattern observed in Clade 2 and Clade 3 (**Fig 4D, 4G, Table S10**). In the case of *bclA2*, pseudogenization events occurred at residue 86 in 79% of cases, all were present in Clade 3. This *bclA2* allele yielded a truncated protein that retained its NTD and the first 72 (17.39%) residues of the CLR (**Fig 4E, 4G, Table S11**). For *bclA3*, pseudogenization events occurred at residue 44, resulting in predicted proteins that retain nearly the full N-terminal domain (NTD, 46 residues), minus two amino acids, all occurring in Clade 3 (**Fig 4F, 4G, Table S12**). These intermediate-length truncations suggest the potential for limited structural function, such as the formation of short, hair-like exosporium projections. Together, these findings indicate that Clade 3 has undergone extensive and consistent *bclA* pseudogenization, likely altering spore surface architecture while preserving partial domain structures in some homologues.

**Fig. 4.**
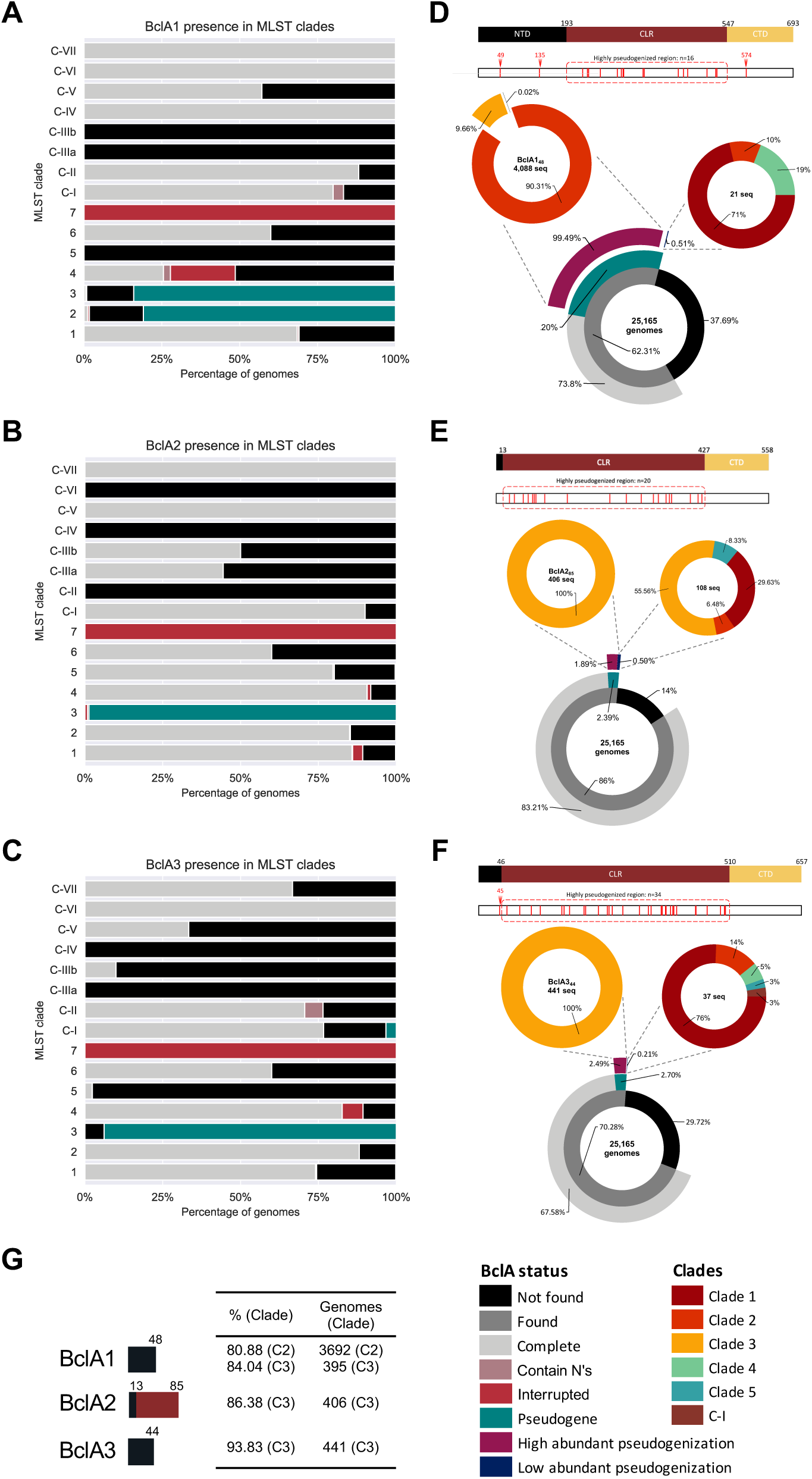
State of BclAs in known diversity of *C. difficile*. Prevalence of found conditions of **(A)** BclA1, **(B)** BclA2 and **(C)** BclA3 on each clade. Prevalence of pseudogenized **(D)** BclA1, **(E)** BclA2 and **(F)** BclA3 sequences per each *C. difficile* clade. Schematic representation of BclAs protein sequences are depicted on top of each donut chart. Truncated BclA sequences were manually curated to locate position of early stop codons, represented as red lines in a white rectangle below protein schematic. **(G)** BclA high abundant pseudogenization alleles present in clade 2 and clade 3. NTD (Black): N-terminal domain, CLR (Maroon): Collagen-like region, CTD (Yellow): C-terminal domain. In this sub-data set, only clades represented in the figure are present in the legend.

The presence of low abundant *bclAs* pseudogenes was also studied. These alleles were pseudogenized in the collagen-like region (red lines in CLR region) (**Fig. 4D, 4E & 4F**), and were distributed among the clades. Since these alleles were unique, it is highly likely that these mutations have no selective pressure associated (**Table S10-S12**).

### Identification and variability of *bclA* alleles

To further identify differences at the amino acid level in the known diversity of *C. difficile* BclAs and determine if there are alleles, a selection of *bclA* genes was used. To do this, deduplicated complete genes were extracted to obtain a set of unique alleles for *bclA1* (1,057, **Table S13**), *bclA2* (1,355, **Table S14**), and *bclA3* (1,842, **Table S15**). For each *bclA* set, sequences were aligned with Clustal Omega against the reference, sequence that presented the least differences compared to all corresponding genes extracted, *bclA1* (CLO_CA1356AA) **(Fig. 5A)**, *bclA2* (CLO_AA3432AA) **(Fig. 5B)** and *bclA3* (CLO_AA0395AA) **(Fig. 5C)**. The alignment was based in codon-aligned amino acids of each BclA set. BclA proteins presented highly conserved NTD and CTD domains with few preserved substitutions events distributed throughout (yellow lines). Remarkably, the CLR region has the most substitutions and Indels occurrence, making it a highly variable region for BclA collagen-like proteins **(Fig. 5).**

**Fig. 5.**
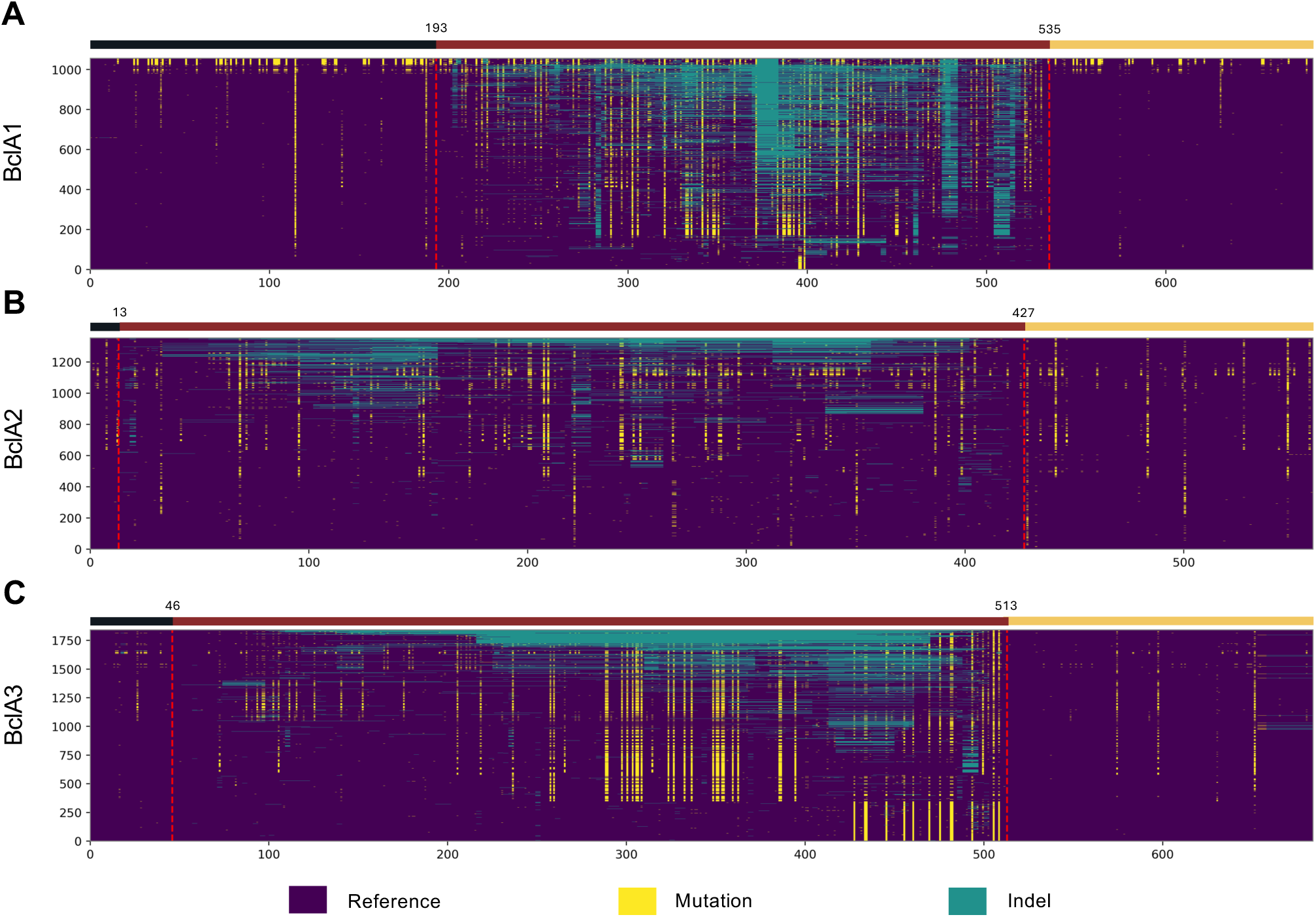
Comparison of *C. difficile* BclA1, BclA2, and BclA3 unique alleles. Unique alleles were aligned using Clustal Omega against the reference. Truncated, interrupted or pseudogenized sequences were not considerer in this analysis. Heatmap displaying positions that differ or not from the reference sequence are depicted for **(A)** BclA1, Ref genome: CLO_CA1356AA, **(B)** BclA2, Ref genome: CLO_AA3432AA and **(C)** BclA3, Ref genome: CLO_AA0395AA. Dashed red lines mark the division between BclA domains. N-terminal domain (NTD, black), Collagen-like region (CLR, maroon) and C-terminal domain (CTD, yellow).

The average amino acid identity was measure for the set of unique BclA alleles. The percentage identity of full length BclA alleles is more than 90% identical in classical clades, but when compared to cryptic clades, a noticeable difference in identity is shown (**Fig S15A, S15B & S15C**). These differences are accentuated when NTD (**Fig. S15D, S15E & S15F**) and CTD (**Fig. S15G, S15H & S15I**) regions are analyzed individually, the identity separation is easily distinguished for all BclA proteins. These findings suggest that the alleles of the classical lineages exhibit a fuzzy separation, and that it is only when compared to the cryptic clades that different types of BclAs become apparent, meaning that classical BclA alleles are very similar. Amino acid identity matrices can be found in Tables S16-S24.

## Discussions

*C. difficile* spores possess an outermost exosporium layer that serves as the initial point of contact with the host. Recent research has revealed the ultrastructural diversity, molecular composition, and functional characteristics of this outer layer ^11, 13–16, 30, 59–61^. Extensive studies have shown that BclA proteins, distinguished by their collagen-like domains, are key contributors to the assembly and structural organization of the exosporium layer in spores of the *Bacillus cereus* group ^31, 34, 62–66^. In contrast, the specific roles and functional implications of BclA-like proteins in *C. difficile* spores remain less clearly defined^14, 16, 17, 52^. Partial characterization in at least two *C. difficile* strains has indicated that these proteins contribute to colonization, interactions with host tissues, spore persistence and recurrence of the disease^14, 16, 17^. In this study, we expanded our insights of the BclA in *C. difficile* by performing a comprehensive analysis of the distribution and variability of the *bclA* orthologues family across over 25,000 *C. difficile* genomes from diverse phylogenetic clades. Furthermore, AlphaFold modeling was employed to structurally characterize all three predicted BclA proteins, identifying key conserved and divergent regions. We also addressed frequent pseudogenization events affecting the *bclA* genes, particularly *bclA1*, and experimentally evaluated the functional consequences by reintroducing a full-length *bclA1* gene into a strain with a pseudogenized version. Our findings offer several major conclusions that collectively advance the understanding of the evolutionary dynamics and functional significance of BclA proteins in *C. difficile* spore biology and pathogenesis.

A major finding of our study is the extensive variability in the presence, sequence conservation, and integrity of *bclA* orthologues across *C. difficile* clades 1 through 5 including the cryptic clades. While *bclA2* and *bclA3* were broadly conserved across most clades, *bclA1* exhibited substantial variability, including complete or partial pseudogenization in multiple lineages—particularly within clade 2 and the cryptic clades. We also observed that clade 5, considered an early ancestor of the four clades, lacked *bclA1* and *bclA3*, while all genomes of clade 3 lack all three *bclA* orthologues. These patterns suggest that selective pressures have driven divergent evolutionary trajectories for spore surface proteins, possibly linked to clade-specific ecological niches or host interactions. It is also tempting to hypothesize that the *bclA* patterns observed across clades might be tied to morphological differences of the hair-like projections. Prior work in strain R20291 that has hair-like projections demonstrates that BclA3 is essential for the formation of the hair-like projections, yet it is unclear whether BclA2 contributes to their formation^16^. Based on these findings, a tentative hypothesis is that all members of clade 3, which lacks all three *bclA* alleles, would be hair-less spores. However, in prior work we demonstrated that spores of clade 1 (strain TL178 [ribotype 002] and TL176 [ribotype 014-020]), clade 2 (strain R20291 [RT027]), and clade 5 (strain M120 [ribotype 078-126]) had hair-like projections, irrespective of the *bclA* encoded genes ^67^. These observations suggest that the formation of hair-like projections on *C. difficile* spores is more complex than previously understood and may involve differential expression of *bclA* orthologues across clades, potentially compensating for the absence or pseudogenization of one or more genes.

The high variability observed in *bclA*s across clades also suggests distinct functional redundancy in virulent traits related to spore host interactions. Functional studies in different clades have shed insight into their roles ^14, 16, 17^. For example, work in clade 1 strain 630, which lacks hair-like projections^11, 17, 30^, shows that *bclA1* is required for proper colonization in mice^17^. However, disruption of individual *bclA*s led to spores lacking the electron-dense exosporium and a defective spore coat, suggesting a pleiotropic effect in the spore surface^17^. By contrast, work on a Clade 2 strain (R20291), shows that a single deletion of *bclA3* is sufficient to impair the formation of the hair-like projection without impacting the formation of the lower electron-dense layers and spore coat^16^. BclA3 was also associated with persistence and disease recurrence^16^. This limited knowledge on *C. difficile* BclAs suggests that depending on the clade, BclAs might have distinct niche-specific advantages, underscoring the need to investigate the impact of BclAs in different clades.

Another major conclusion offered by this work is the striking pattern of sequence conservation and variability across the BclA proteins-BclA1, BclA2 and BclA3. In each case, both the N-terminal and C-terminal domains were highly conserved across the more than 25,000 *bclA* alleles analyzed, while the central collagen-like regions exhibited substantial sequence diversity. This fact that conserved terminal domains flank a highly variable central region, suggests preservation of the domains requires for BclA anchoring and structural stability (the NTD and CTD, respectively), while allowing evolutionary flexibility in the central collagen-like region. The pronounced variability in the collagen-like domains of all three BclA may be attributed to the repetitive GXY triplet motif that results in underlying DNA sequence repeats. Such repeats are known to increase genetic instability through replication slippage or unequal recombination, which can result in insertions, deletions, or sequence rearrangements^68^. While we did not investigate further in this work, it is likely that in *C. difficile* the collagen-like domain may be subject to diversification to host-derived selective pressures, including evasion host immune responses or adapt to distinct host environments as observed in *B. anthracis* spores^69, 70^. Together, these findings indicate that the collagen-like region is a shared hotspot for adaptive variation across the BclA protein family in *C. difficile*. Additionally, these observations deserve further work to address how variations in the collagen-like region affects spore adherence, environmental persistence, or host colonization dynamics.

Another notable finding from our analysis is the pattern of *bclA1* pseudogenization across *C. difficile* clades. While pseudogenization events in *bclA* genes were generally rare, occurring at a frequency below 1% across most clades, a striking exception was observed in Clade 2. In this clade, most strains carried a conserved pseudogenized *bclA1* allele that encodes only a short 48-amino acid peptide corresponding to the N-terminal domain (NTD) of BclA1. Despite the truncation, this NTD fragment appears to retain the structural features necessary for anchoring to the spore surface^17^. The conservation of this pseudogenized form within Clade 2 is particularly intriguing, as it suggests that full-length BclA1 may be dispensable in this lineage, or that the truncated form fulfills a distinct, potentially clade-specific function. This raises questions about the evolutionary pressures shaping *bclA1* function and whether the presence of only the anchoring domain represents an adaptive modification.

In summary, this work highlights the diversity of the collagen-like BclA proteins in *C. difficile* and demonstrate that they exhibit conserved modular domains defined by a highly conserved N- and C-terminal domains, flanked by a highly variable collagen-like region. While the precise implications of Clade-specific variabilities remains unclear,

## Supplementary figure legend

**Fig. S1. *C. difficile bclA1* gene schematic representation in 630 & R20291 strain.** *bclA1_630_* (CD630_03320) is a 2082 bp gene coding a 694 amino acid protein (67.8 kDa). *bclA1_R20291_* (CDIF27147_00471, CDIF27147_00472, CDIF27147_00473) is represented as a three-segmented Open Reading Frame (ORF) (*bclA1_1, bclA1_2 & bclA1_3*) because it is a pseudogenized gene. A nonsense mutation, A145T, generated an early stop codon resulting in a short protein of 48 amino acids (4.7 kDa). Downstream, another nonsense mutation, C739T, also generates an early stop codon and the two extra segments of *bclA1_R20291_* ORF*. bclA1* possess a σ^K^ promoter region (Saujet, 2013), and two putative σ^E^ and σ^G^ consensus sequences that were found during this work. *bclA1* is flanked upstream by *ppiB* and downstream by *ppaC*. *C. difficile* 630 genome used as reference was GenBank ID: AM180355.1. *C. difficile* R20291 genome used as reference was GenBank ID: CP029423. Scale bar: 1 kb.

**Fig. S2. BclA1 Pairwise sequence alignment (PSA) of 630 & R20291 strains.** EMBOSS Needle was used for PSA. A representation of the protein domains is depicted above the sequence. The proteins possess a 51.4% Identity. Repaired BclA1 from R20291 was used for the alignment. N-terminal domain (NTD, Black), Collagen-like region (CLR, Maroon) and C-terminal domain (CTD, Yellow). *C. difficile* 630 genome used as reference was GenBank ID: AM180355.1. *C. difficile* R20291 genome used as reference was GenBank ID: CP029423.

**Fig. S3. *C. difficile bclA2* gene schematic representation in 630 & R20291 strains.** *bclA2_630_* (CD630_32300) is a 1677 bp gene coding a 558 amino acid protein (49.1 kDa). *bclA2_R20291_* (CDIF27147_03409) is a 1641 bp gene coding a 546 amino acid protein (47.9 kDa). *bclA2* possess a σ^K^ promoter region (Saujet, 2013) and a putative σ^E^ consensus sequence that was found during this work. *bclA2* is flanked upstream by *hpt2* and downstream by *dapB2*. *C. difficile* 630 genome used as reference was GenBank ID: AM180355.1. *C. difficile* R20291 genome used as reference was GenBank ID: CP029423. Scale bar: 1 kb.

**Fig. S4. BclA2 Pairwise sequence alignment (PSA) of 630 & R20291 strains.** EMBOSS Needle was used for PSA. A representation of the protein domains is depicted above the sequence. The proteins possess a 95.2% Identity. N-terminal domain (NTD, Black), Collagen-like region (CLR, Maroon) and C-terminal domain (CTD, Yellow). *C. difficile* 630 genome used as reference was GenBank ID: AM180355.1. *C. difficile* R20291 genome used as reference was GenBank ID: CP029423.

**Fig. S5. *C. difficile bclA3* gene schematic representation in 630 & R20291 strains.** *bclA3_630_* (CD630_33490) is a 1986 bp gene coding a 661 amino acid protein (58.3 kDa). *bclA3_R20291_* (CDIF27147_03519) is a 2037 bp gene coding a 678 amino acid protein (59.9 kDa). *bclA3* is the second gene of an operon with an upstream *sgtA* gene coding for a glycosyl transferase (CD630_33500, CDIF27147_03520). The operon possesses a σ^K^ promoter region, and an additional σ^K^ promoter region (Saujet, 2013) immediately upstream *bclA3* and a putative σ^E^ consensus sequence that was found upstream the operon during this work. *bclA3* is flanked upstream by *sgtA* and downstream by CDIF27147_03518. *C. difficile* 630 genome used as reference was GenBank ID: AM180355.1. *C. difficile* R20291 genome used as reference was GenBank ID: CP029423. Scale bar: 1 kb.

**Fig. S6. BclA3 Pairwise sequence alignment (PSA) of 630 & R20291 strains.** EMBOSS Needle was used for PSA. A representation of the protein domains is depicted above the sequence. The proteins possess an 86.9% Identity. N-terminal domain (NTD, Black), Collagen-like region (CLR, Maroon) and C-terminal domain (CTD, Yellow). *C. difficile* 630 genome used as reference was GenBank ID: AM180355.1. *C. difficile* R20291 genome used as reference was GenBank ID: CP029423.

**Fig. S7. Full length BclA1 Alphafold3 predicted structure.** Predicted monomeric structure of *C. difficile* **(A)** BclA1_630_, **(B)** rBclA1_R20291_, and predicted homotrimer structures of **(C)** BclA1_630_ and **(D)** rBclA1_R20291_. Each monomer unit has a different color for easy recognition in trimeric structures. Each model has its respective Predicted aligned error (PAE) plot below. NTD: N-terminal domain, CLR: Collagen-like region, CTD: C-terminal domain.

**Fig. S8. Full length BclA2_R20291_ Alphafold3 predicted structure.** Predicted structure of *C. difficile* **(A)** BclA2_R20291_ monomer and **(B)** homotrimer complex. Each monomer unit has a different color for easy recognition in trimeric structures. Each model has its respective Predicted aligned error (PAE) plot below. NTD: N-terminal domain, CLR: Collagen-like region, CTD: C-terminal domain.

**Fig. S9. Full length BclA3_R20291_ Alphafold3 predicted structure.** Predicted structure of *C. difficile* **(A)** BclA3_R20291_ monomer and **(B)** homotrimer complex. Each monomer unit has a different color for easy recognition in trimeric structures. Each model has its respective Predicted aligned error (PAE) plot below. NTD: N-terminal domain, CLR: Collagen-like region, CTD: C-terminal domain.

**Fig. S10. AlphaFold3 predicted structure of BclA_10GXY homotrimer.** Predicted structure of *C. difficile* **(A)** BclA1_630_, **(B)** BclA2_R20291_ and **(C)** BclA3_R20291_ homotrimer complexes. Each model has its respective Predicted aligned error (PAE) plot below.

**Fig. S11. AlphaFold3 predicted CTD heterotrimer structures from R20291 strain.** Heterotrimer predicted model composed of **(A)** 2 units of BclA2_CTD and 1 unit of BclA3_CTD or **(B)** 1 unit of BclA2_CTD and 2 unit of BclA3_CTD. Each model has its respective Predicted aligned error (PAE) plot below.

**Fig. S12. AlphaFold3 predicted CTD heterotrimer structures from 630 strain.** Heterotrimer prediction composed of **(A)** 2 units of BclA1_CTD and 1 unit of BclA2_CTD, **(B)** 2 units of BclA1_CTD and 1 unit of BclA3_CTD, **(C)** 1 unit of BclA1_CTD and 2 unit of BclA2_CTD, **(D)** 1 unit of BclA1_CTD and 2 unit of BclA3_CTD or **(E)** 1 unit of BclA1_CTD, 1 unit of BclA2_CTD and 1 unit of BclA3_CTD. Each model has its respective Predicted aligned error (PAE) plot.

**Fig. S13. Pairwise alignment (PSA) of *C. difficile* BclA_CTD with *B. anthracis* BclA/BclB_CTD.** EMBOSS Needle was used for PSA of **(A)** Cd_BclA1_CTD_630_ with Ba_BclA_CTD **(B)** Cd_BclA2_CTD_R20291_ with Ba_BclA_CTD and **(C)** Cd_BclA3_CTD_R20291_ with Ba_BclB_CTD. **(D)** Percentage of identity of aligned sequences. CTD: C-terminal domain. *C. difficile* 630 genome used as reference was GenBank sequence: AM180355.1. *C. difficile* R20291 genome used as reference was GenBank sequence: CP029423. *B. anthracis* genome used as reference was GenBank sequence: AE016879.1.

**Fig. S14. Distribution of BclAs in known diversity of *C. difficile*. (A)** Zoomed-in view of the Clade 4 subtree. **(B)** Zoomed-in view of the Clade 5 subtree. **(C)** Zoomed-in view of the cryptic clades subtree. Horizontal bars represent the state of BclA in a particular genome (not found, complete, containing Ns, interrupted, or pseudogenized).

**Fig. S15. Average amino acid identity (AAI) analysis of BclA unique alleles.** Heatmap of the percentage of identity between all unique alleles of **(A)** BclA1, **(B)** BclA2 and **(C)** BclA3. AAI was performed for full length protein, NTD and CTD regions. N-terminal domain (NTD) and C-terminal domain (CTD).

