## Supplementary Figures for "Phylogenomic analysis of the collagen-like BclA proteins in *Clostridioides difficile*"

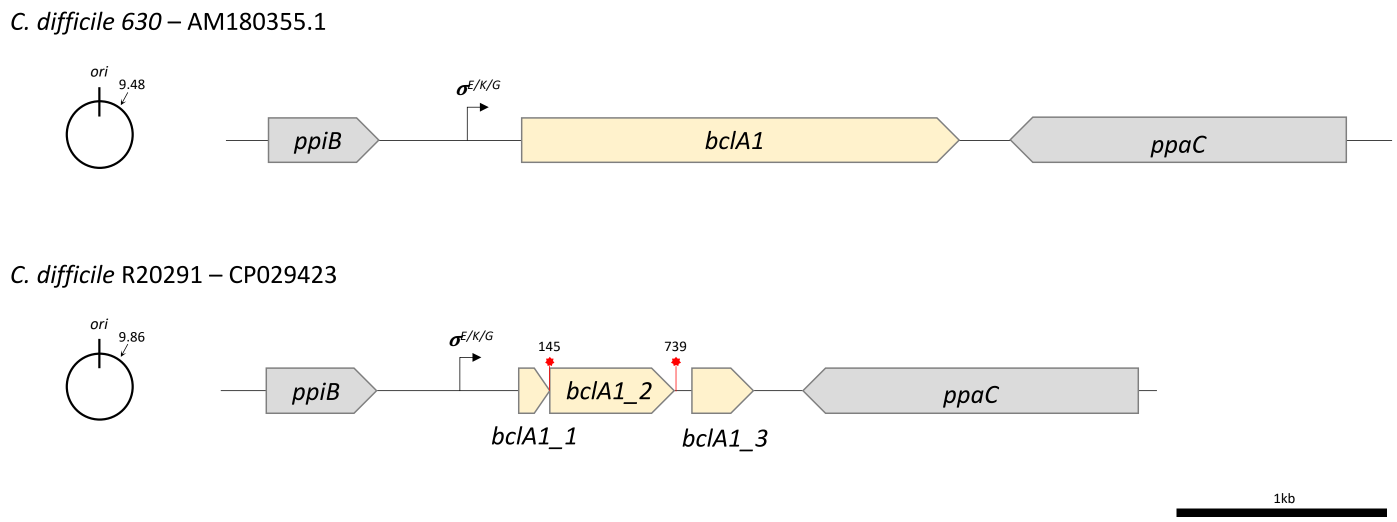


**Fig. S1. *C. difficile bclA1* gene schematic representation in 630 & R20291 strain**. $bclA1_{630}$ (CD630_03320) is a 2082 bp gene coding a 694 amino acid protein (67.8 kDa). $bclA1_{R20291}$ (CDIF27147_00471, CDIF27147_00472, CDIF27147_00473) is represented as a three-segmented Open Reading Frame (ORF) (*bclA1_1, bclA1_2* & *bclA1_3*) because it is a pseudogenized gene. A nonsense mutation, A145T, generated an early stop codon resulting in a short protein of 48 amino acids (4.7 kDa). Downstream, another nonsense mutation, C739T, also generates an early stop codon and the two extra segments of $bclA1_{R20291}$ ORF. *bclA1* possess a $\sigma^K$ promoter region (Saujet, 2013), and also two putative $\sigma^E$ and $\sigma^G$ consensus sequences that were found during this work. *bclA1* is flanked upstream by *ppiB* and downstream by *ppaC*. *C. difficile* 630 genome used as reference was GenBank ID: AM180355.1. *C. difficile* R20291 genome used as reference was GenBank ID: CP029423. Scale bar: 1 kb.

```
                                             NTD
BclA1_630        1 MRNIILYLNDDTFISKKYPDKNFSNLDYCLIGSKCSNSFVKEKLITFFKV     50
                   ||.|||||||||||||||||||||||||||||||||||||||||||||||
BclA1_R20291     1 MRKIILYLNDDTFISKKYPDKNFSNLDYCLIGSKCSNSFVKEKLITFFKV     50

BclA1_630       51 RIPDILKDKSILKAELFIHIDSNKNHIFKEKVDIEIKRISEYYNLRTITW    100
                   ||||||||||||||||||||||||||||||||||||||||||||||||||
BclA1_R20291    51 RIPDILKDKSILKAELFIHIDSNKNHIFKEKVDIEIKRISEYYNLRTITW    100

BclA1_630      101 NDRVSMENIRGYLPIGISDTSNYICLNITGTIKAWAMNKYPNYGLALSLN    150
                   ||||||||||||||||||||||||||||||||||||||||||||||||||
BclA1_R20291   101 NDRVSMENIRGYLPIGISDTSNYICLNITGTIKAWAMNKYPNYGLALSLN    150

BclA1_630      151 YPYQILEFTSSRGCNKPYILVTFEDRIIDNCYPKCECPPIRITGPMGPRG    200
                   |||||.||||||.||||||||||||||||||||||||.||||||||||||
BclA1_R20291   151 YPYQIFEFTSSRDCNKPYILVTFEDRIIDNCYPKCECLPIRITGPMGPRG    200

                                            CLR
BclA1_630      201 ATGSTGPMGVTGPTGSTGATGSIGPTGPTGNTGATGSIGPTGVTGPTGST    250
                   ||||.||||.
BclA1_R20291   201 ATGSIGPMGA----------------------------------------    210

BclA1_630      251 GATGSIGPTGVTGPTGNTGVTGSIGPTGATGPTGNTGVTGSIGPTGVTGP    300

BclA1_R20291   210 --------------------------------------------------    210

BclA1_630      301 TGNTGEIGPTGATGPTGVTGSIGPTGATGPTGEIGPTGATGATGSIGPTG    350

BclA1_R20291   210 --------------------------------------------------    210

BclA1_630      351 ATGPTGATGVTGEIGPTGEIGPTGATGPTGVTGSIGPTGATGPTGATGEI    400

BclA1_R20291   210 --------------------------------------------------    210

BclA1_630      401 GPTGATGPTGVTGSIGPTGATGPTGATGEIGPTGATGPTGVTGEIGPTGA    450

BclA1_R20291   210 --------------------------------------------------    210

BclA1_630      451 TGPTGNTGVTGEIGPTGATGPTGNTGVTGEIGPTGATGPTGVTGEIGPTG    500

BclA1_R20291   210 --------------------------------------------------    210

BclA1_630      501 NTGATGSIGPTGVTGPTGATGSIGPTGATGATGVTGPTGPTGATGNSSQP    550
                                                        |||||||||||||
BclA1_R20291   211 -------------------------------------TGPTGATGNSSQP    223

                                            CTD
BclA1_630      551 VANFLVNAPSPQTLNNGDAITGWQTIIGNSSSITVDTNGTFTVQENGVYY    600
                   :||||||||||||||||:||||||:|||||||||||.|||||||||||||
BclA1_R20291   224 IANFLVNAPSPQTLNNGNAITGWKTIIGNSSSITVDANGTFTVQENGVYY    273

BclA1_630      601 ISVSVALQPGSSSINQYSFAILFPILGGKDLAGLTTEPGGGGVLSGYFAG    650
                   ||||||||||||||||||||||||||||||||||||||||||||||||||
BclA1_R20291   274 ISVSVALQPGSSSINQYSFAILFPILGGKDLAGLTTEPGGGGVLSGYFAG    323

BclA1_630      651 FLFGGTTFTINNFSSTTVGIRNGQSAGTAATLTIFRIADTVMT    693
                   |||||||||||||||||||||||||||||||||||||||||||
BclA1_R20291   324 FLFGGTTFTINNFSSTTVGIRNGQSAGTAATLTIFRIADTVMT    366
```

**Fig. S2. BclA1 Pairwise sequence alignment (PSA) of 630 & R20291 strains**. EMBOSS Needle was used for PSA. A representation of the protein domains is depicted above the sequence. The proteins possess a **51.4%** Identity. Repaired BclA1 from R20291 was used for the alignment. N-terminal domain (NTD, Black), Collagen-like region (CLR, Maroon) and C-terminal domain (CTD, Yellow). *C. difficile* 630 genome used as reference was GenBank ID: AM180355.1. *C. difficile* R20291 genome used as reference was GenBank ID: CP029423.

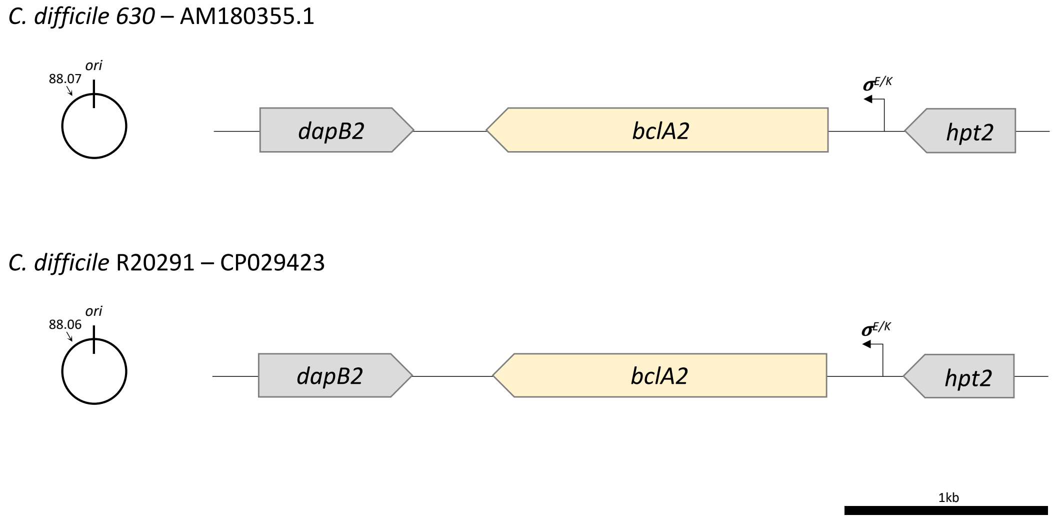


**Fig. S3. *C. difficile bclA2* gene schematic representation in 630 & R20291 strains**. *bclA2*$_{630}$ (CD630_32300) is a 1677 bp gene coding a 558 amino acid protein (49.1 kDa). *bclA2*$_{R20291}$ (CDIF27147_03409) is a 1641 bp gene coding a 546 amino acid protein (47.9 kDa). *bclA2* possess a $\sigma^K$ promoter region (Saujet, 2013) and a putative $\sigma^E$ consensus sequence that was found during this work. *bclA2* is flanked upstream by *hpt2* and downstream by *dapB2*. *C. difficile* 630 genome used as reference was GenBank ID: AM180355.1. *C. difficile* R20291 genome used as reference was GenBank ID: CP029423. Scale bar: 1 kb.

```
                       NTD
BclA2_630          1 MSDISGPSLYQDVGPTGPTGATGPTGPTGPRGATGATGANGITGPTGNTG     50
                     ||||||||||||||||||||||||||||||||||||||||||||||||||
BclA2_R20291       1 MSDISGPSLYQDVGPTGPTGATGPTGPTGPRGATGATGANGITGPTGNTG     50

BclA2_630         51 ATGANGITGPTGNMGATGPNGTTGSTGPTGNTGATGANGITGPTGNTGAT    100
                     ||||||||||||||||||.||||||||||||||||||||||||||.||||
BclA2_R20291      51 ATGANGITGPTGNMGATGANGTTGSTGPTGNTGATGANGITGPTGATGAT    100

BclA2_630        101 GANGITGPTGNKGATGANGITGSTGPTGNTGATGANGITGPTGNTGATGA    150
                     ||||||||||||||||||||   |||||.|||||||||||||||||||||
BclA2_R20291     101 GANGITGPTGNKGATGANGI---TGPTGATGATGANGITGPTGNTGATGA    147

BclA2_630        151 TGPTGLTGATGATGANGITGPTGNTGATGANGVTGATGPTGNTGATGPTG    200
                     .|.||||||||||||||||||||.||||||||||||||||||||||||||
BclA2_R20291     148 NGATGLTGATGATGANGITGPTGATGATGANGVTGATGPTGNTGATGPTG    197

                                                CLR
BclA2_630        201 SIGATGATGTTGATGPIGATGATGADGEVGPTGAVGATGPDGLVGPTGPT    250
                     |||||||.|.||||||||||         |||||||||||||||||||||
BclA2_R20291     198 SIGATGANGVTGATGPIGAT---------GPTGAVGATGPDGLVGPTGPT    238

BclA2_630        251 GPTGATGANGLVGPTGPTGATGANGLVGPTGATGATGVAGAIGPTGAVGA    300
                     ||||||||||||||||||||||||||||||||||||||||||||||||||
BclA2_R20291     239 GPTGATGANGLVGPTGPTGATGANGLVGPTGATGATGVAGAIGPTGAVGA    288

BclA2_630        301 TGPTGADGAVGPTGATGATGANGATGPTGAVGATGANGVAGPIGPTGPTG    350
                     ||||||||||||||||||||||||||||||||||||||||||||||||||
BclA2_R20291     289 TGPTGADGAVGPTGATGATGANGATGPTGAVGATGANGVAGPIGPTGPTG    338

BclA2_630        351 ENGVAGATGATGATGANGATGPTGAVGATGANGVAGAIGPTGPTGANGAT    400
                     .|||||||||||||||||||||||||||||||||||.|||||||||||.|
BclA2_R20291     339 ANGVAGATGATGATGANGATGPTGAVGATGANGVAGPIGPTGPTGANGTT    388

                                                          CTD
BclA2_630        401 GATGATGATGANGATGPTGATGATGVLAANNAQFTVSSSSLVNNTLVTFN    450
                     |||||||||||||||||||||||||||||||||||||||||.||||||||
BclA2_R20291     389 GATGATGATGANGATGPTGATGATGVLAANNAQFTVSSSSLGNNTLVTFN    438

BclA2_630        451 SSFINGTNITFPTSSTINLAVGGIYNVSFGIRATLSLAGFMSITTNFNGV    500
                     |||||||||||||||||||||||||||||||||.||||||||||||||||
BclA2_R20291     439 SSFINGTNITFPTSSTINLAVGGIYNVSFGIRAILSLAGFMSITTNFNGV    488

BclA2_630        501 TQNNFIAKAVNTLTSSDVSVSLSFLVDARAAAVTLSFTFGSGTTGTSAAG    550
                     .||||||||||||||||||||||||||||||||||||||||||||||.||
BclA2_R20291     489 AQNNFIAKAVNTLTSSDVSVSLSFLVDARAAAVTLSFTFGSGTTGTSPAG    538

BclA2_630        551 YVSVYRIQ    558
                     ||||||||
BclA2_R20291     539 YVSVYRIQ    546
```

**Fig. S4. BclA2 Pairwise sequence alignment (PSA) of 630 & R20291 strains**. EMBOSS Needle was used for PSA. A representation of the protein domains is depicted above the sequence. The proteins possess a **95.2%** Identity. N-terminal domain (NTD, Black), Collagen-like region (CLR, Maroon) and C-terminal domain (CTD, Yellow). *C. difficile* 630 genome used as reference was GenBank ID: AM180355.1. *C. difficile* R20291 genome used as reference was GenBank ID: CP029423.

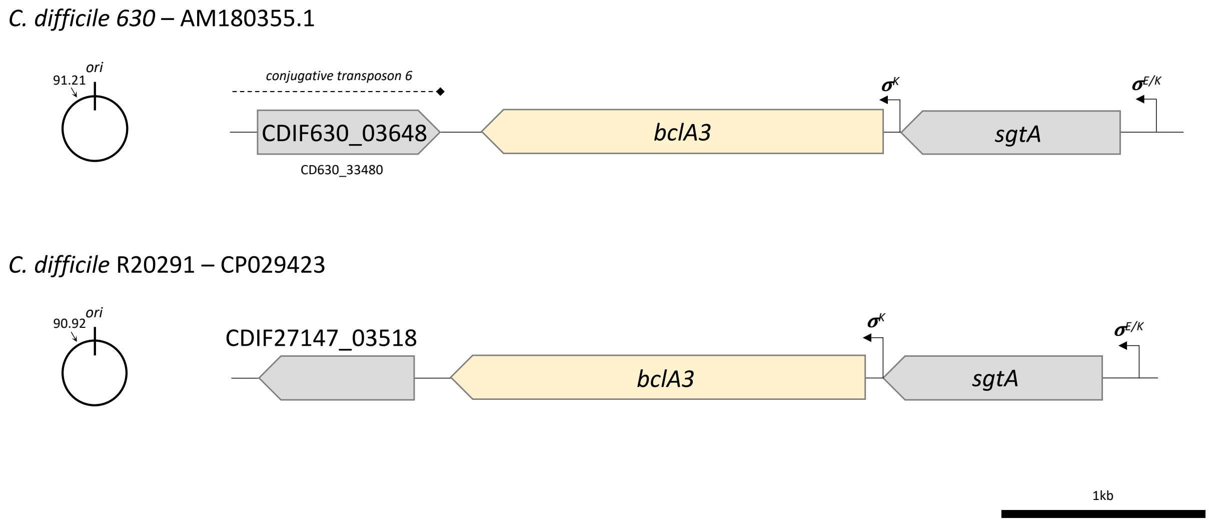


**Fig. S5. *C. difficile bclA3* gene schematic representation in 630 & R20291 strains**. *bclA3*$_{630}$ (CD630_33490) is a 1986 bp gene coding a 661 amino acid protein (58.3 kDa). *bclA3*$_{R20291}$ (CDIF27147_03519) is a 2037 bp gene coding a 678 amino acid protein (59.9 kDa). *bclA3* is the second gene of an operon with an upstream *sgtA* gene coding for a glycosyl transferase (CD630_33500, CDIF27147_03520). The operon posses a $\sigma^K$ promoter region, and an additional $\sigma^K$ promoter region (Saujet, 2013) immediately upstream *bclA3* and also a putative $\sigma^E$ consensus sequence that was found upstream the operon during this work. *bclA3* is flanked upstream by *sgtA* and downstream by CDIF27147_03518. *C. difficile* 630 genome used as reference was GenBank ID: AM180355.1. *C. difficile* R20291 genome used as reference was GenBank ID: CP029423. Scale bar: 1 kb.

```
                                  NTD
BclA3_630          1 MSRNKYFGPFDDNDYNNGYDKYDDCNNGRDDYNSCDCHHCCPPSCVGPTG     50
                     ||||||||||||||||||||||||||||||||||||||||||||||||||
BclA3_R20291       1 MSRNKYFGPFDDNDYNNGYDKYDDCNNGRDDYNSCDCHHCCPPSCVGPTG     50

BclA3_630         51 PMGPRGRTGPTGPTGPTGPGVGATGPTGPTGPTGPTGNTGNTGATGLRGP    100
                     ||||||||||||||||||||||.|||||||||||||||||||||||||||
BclA3_R20291      51 PMGPRGRTGPTGPTGPTGPGVGGTGPTGPTGPTGPTGNTGNTGATGLRGP    100

BclA3_630        101 TGATGATGPTGATGAIGFGV---TGPTGPTGATGATGADGVTGPTGPTGA    147
                     |||||.||||||||||||||   |||||||||||||||||||||||||||
BclA3_R20291     101 TGATGGTGPTGATGAIGFGVTGPTGPTGPTGATGATGADGVTGPTGPTGA    150

BclA3_630        148 TGADGITGPTGATGATGFGVTGPTGPTGATGVGVTGATGLIGPTGATGTP    197
                     ||||||||||||||||||||||||||||||||||||||||||||||||||
BclA3_R20291     151 TGADGITGPTGATGATGFGVTGPTGPTGATGVGVTGATGLIGPTGATGTP    200

                                           CLR
BclA3_630        198 GATGPTGAIGATGIGITGPTGATGATGADGATGVTGPTGPTGATGADGVT    247
                     ||||||||||||||||||||||||||||||||||||||||||||||||||
BclA3_R20291     201 GATGPTGAIGATGIGITGPTGATGATGADGATGVTGPTGPTGATGADGVT    250

BclA3_630        248 GPTGATGATGIGITGPTGPTGATGIGITGATGLIGPTGATGTPGATGPTG    297
                     ||||||||||||||||||.||||||||||||||||||||||..|||||||
BclA3_R20291     251 GPTGATGATGIGITGPTGATGATGIGITGATGLIGPTGATGATGATGPTG    300

BclA3_630        298 ATGPTG-VGVTGATGATGATGADGATGVTGPTGATGATGANGLVGPTGAT    346
                     .||.|| .|:.|.|||||.||||||||.||.|||||.|||:|||||||||
BclA3_R20291     301 VTGATGAAGLIGPTGATGVTGADGATGATGATGATGPTGADGLVGPTGAT    350

BclA3_630        347 GAAGTPGATGPTGATGPTGVGITGATGATGATGPTGADGATGPTGATGNT    396
                     ||.|..|..||||.||.||||||||||||||||||||||..|||||||.|
BclA3_R20291     351 GATGADGLVGPTGPTGATGVGITGATGATGATGPTGADGLVGPTGATGAT    400

BclA3_630        397 GADGVAGP---TGATGNTGADGATGPTGATGATGADGATGPTGATGATGV    443
                     ||||||||   ||||||||||||||||||||.|||||..||||||||||:
BclA3_R20291     401 GADGVAGPTGATGATGNTGADGATGPTGATGPTGADGLVGPTGATGATGL    450

BclA3_630        444 AGATGATGP---TGATGADGATGPTGATGATGADGATGPTGATGATGVTG    490
                     |||||||||   ||.||||||||.|||||.|||||..||||||||   ||
BclA3_R20291     451 AGATGATGPIGATGPTGADGATGATGATGPTGADGLVGPTGATGA---TG    497

BclA3_630        491 ATGPTGPTGATGATGATGASAIIPFASGIPLSLTTIAGGLVGTPGFVGFG    540
                     |||||||         ||||||||||||||||||||||||||||||||||
BclA3_R20291     498 ATGPTGP---------TGASAIIPFASGIPLSLTTIAGGLVGTPGFVGFG    538

                                           CTD
BclA3_630        541 SSAPGLSIVGGVIDLTNAAGTLTNFAFSMPRDGTITSISAYFSTTAALSL    590
                     ||||||||||||||||||||||||||||||||||||||||||||||||||
BclA3_R20291     539 SSAPGLSIVGGVIDLTNAAGTLTNFAFSMPRDGTITSISAYFSTTAALSL    588

BclA3_630        591 VGSTITITATLYQSTAPNNSFTAVPGATVTLAPPLTGILSVGSISSGIVT    640
                     ||||||||||||||||||||||||||||||||||||||||||||||||||
BclA3_R20291     589 VGSTITITATLYQSTAPNNSFTAVPGATVTLAPPLTGILSVGSISSGIVT    638

BclA3_630        641 GLNIAATAQTPDRQYAI--------------------------    657
                     ||||||||:|   ::.:
BclA3_R20291     639 GLNIAATAET---RFLLVFTATASGLSLVNTVAGYASAGIAIN    678
```

**Fig. S6. BclA3 Pairwise sequence alignment (PSA) of 630 & R20291 strains**. EMBOSS Needle was used for PSA. A representation of the protein domains is depicted above the sequence. The proteins possess an **86.9%** Identity. N-terminal domain (NTD, Black), Collagen-like region (CLR, Maroon) and C-terminal domain (CTD, Yellow). *C. difficile* 630 genome used as reference was GenBank ID: AM180355.1. *C. difficile* R20291 genome used as reference was GenBank ID: CP029423.

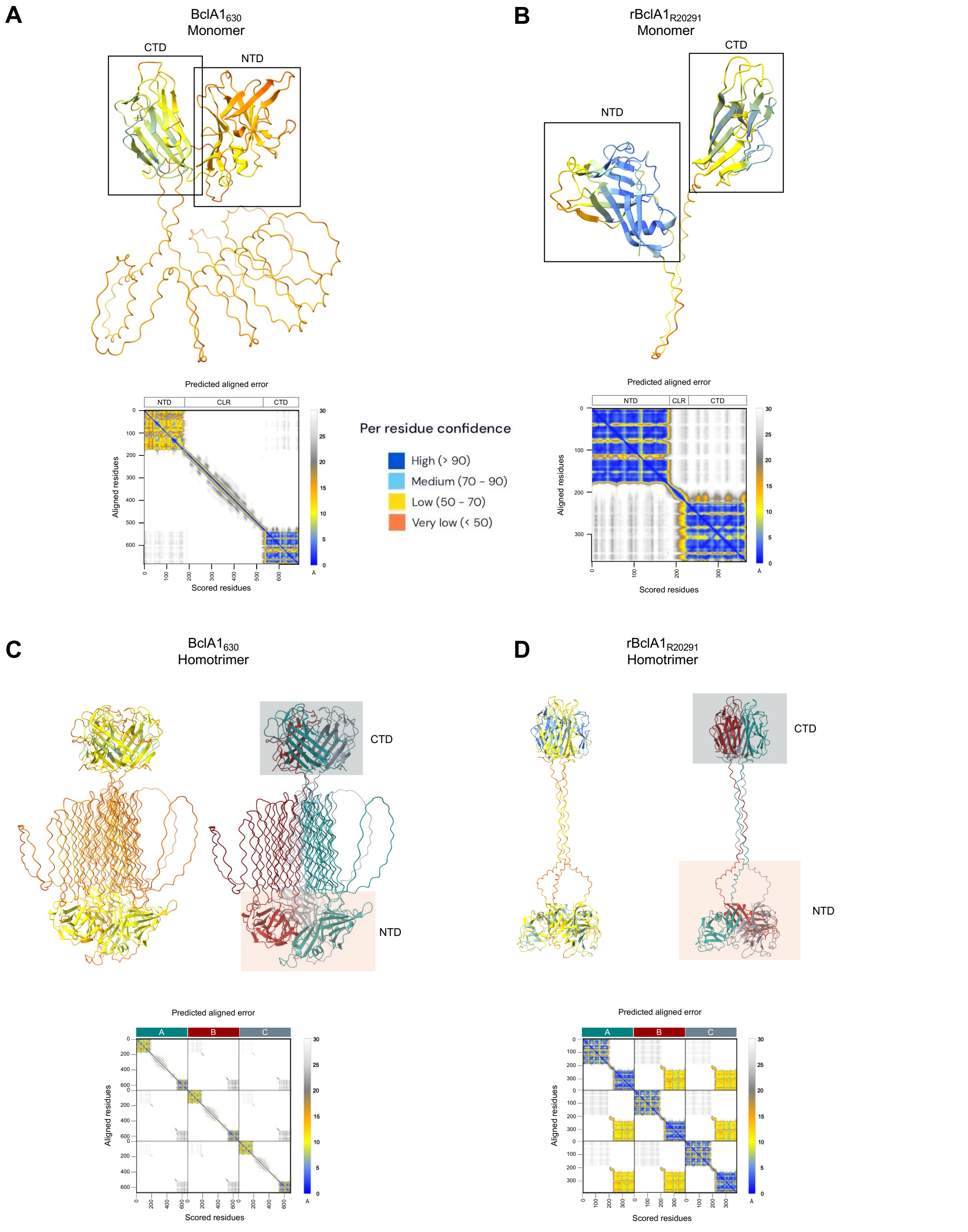


**Fig. S7. Full length BclA1 Alphafold3 predicted structure.** Predicted monomeric structure of *C. difficile* **(A)** BclA1$_{630}$, **(B)** rBclA1$_{R20291}$, and predicted homotrimer structures of **(C)** BclA1$_{630}$ and **(D)** rBclA1$_{R20291}$. Each monomer unit has a different color for easy recognition in trimeric structures. Each model has its respective Predicted aligned error (PAE) plot below. NTD: N-terminal domain, CLR: Collagen-like region, CTD: C-terminal domain.

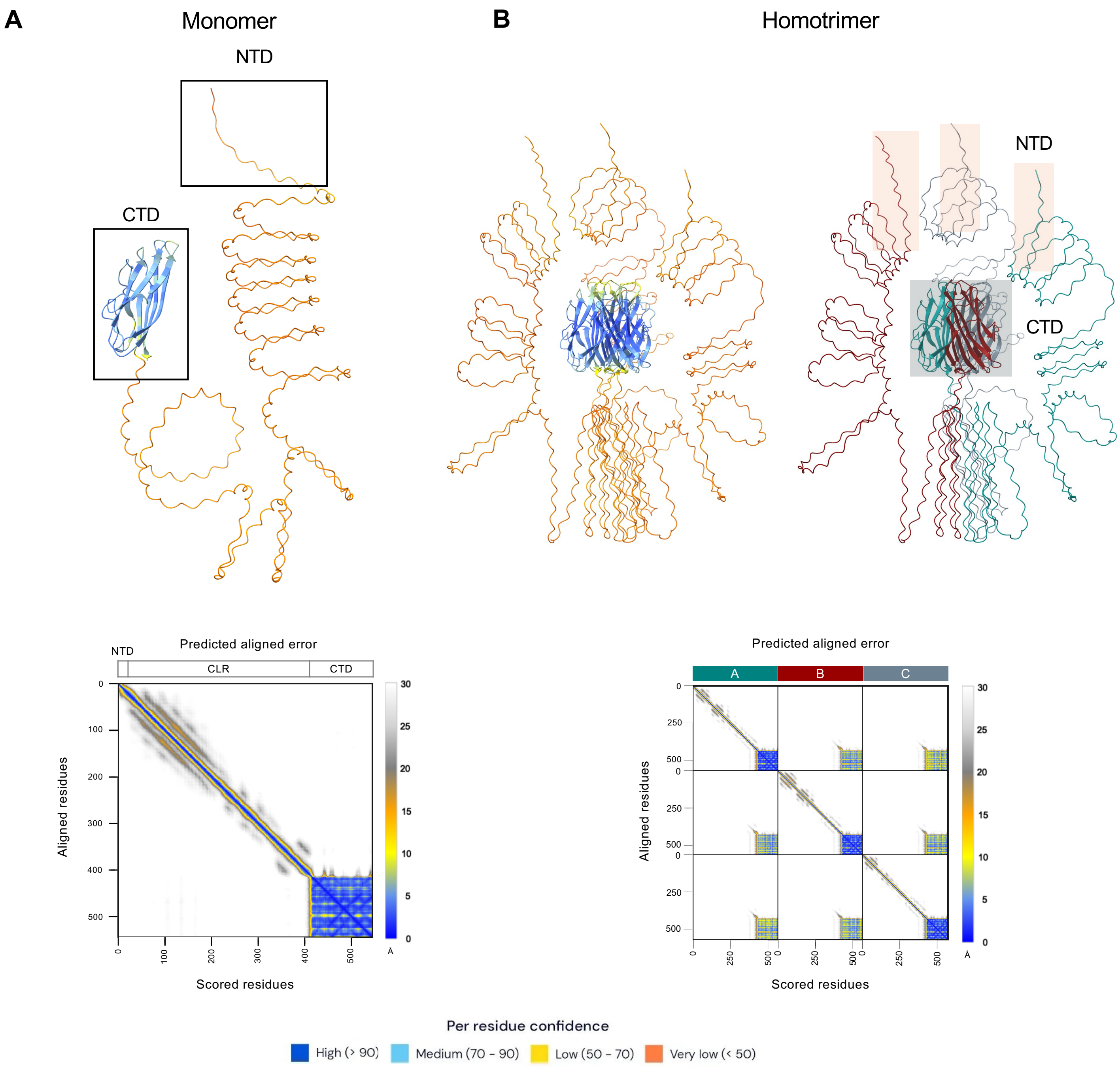


**Fig. S8. Full length BclA2$_{R20291}$ Alphafold3 predicted structure.** Predicted structure of *C. difficile* **(A)** BclA2$_{R20291}$ monomer and **(B)** homotrimer complex. Each monomer unit has a different color for easy recognition in trimeric structures. Each model has its respective Predicted aligned error (PAE) plot below. NTD: N-terminal domain, CLR: Collagen-like region, CTD: C-terminal domain.

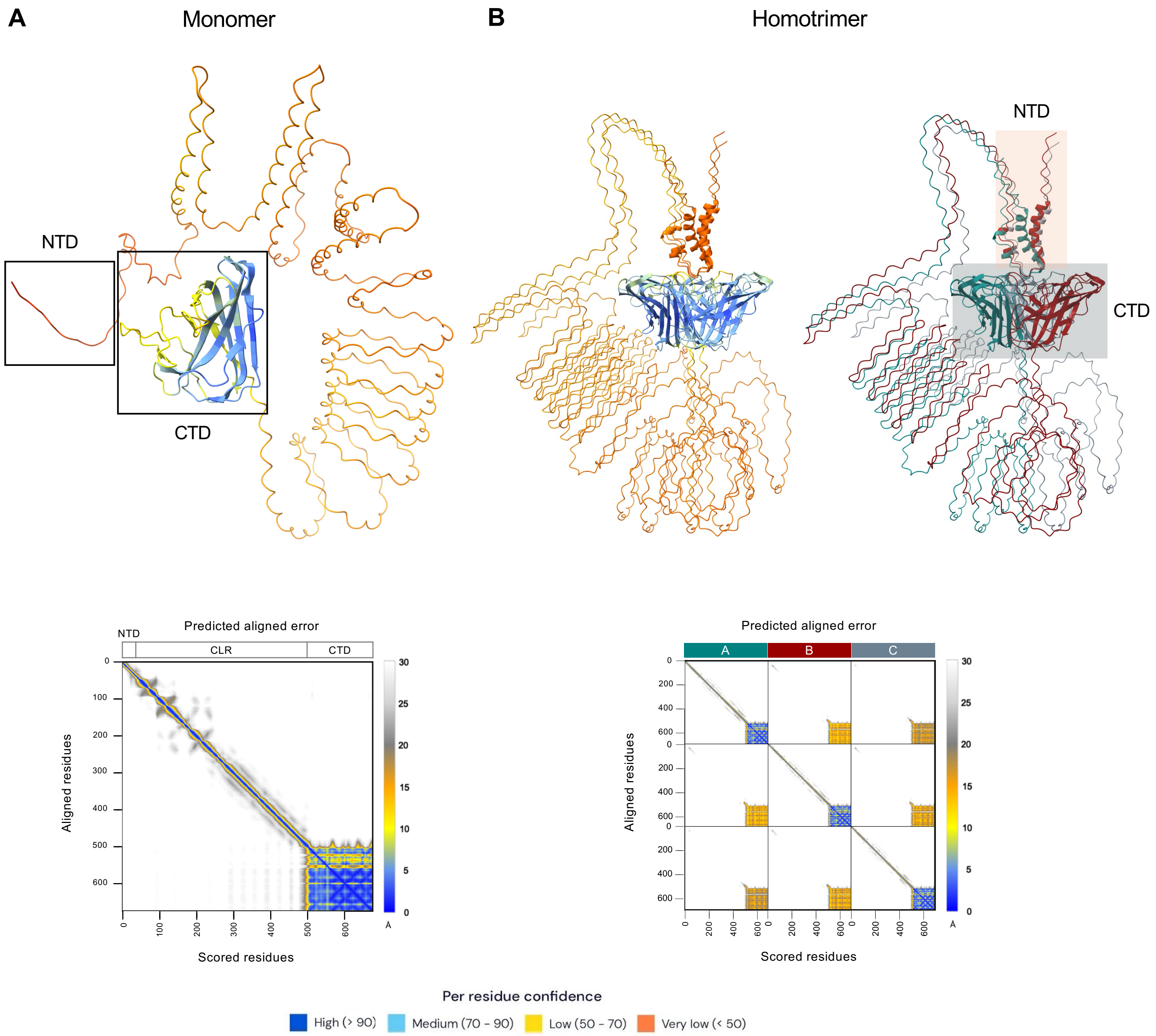


**Fig. S9. Full length BclA3$_{R20291}$ Alphafold3 predicted structure.** Predicted structure of *C. difficile* **(A)** BclA3$_{R20291}$ monomer and **(B)** homotrimer complex. Each monomer unit has a different color for easy recognition in trimeric structures. Each model has its respective Predicted aligned error (PAE) plot below. NTD: N-terminal domain, CLR: Collagen-like region, CTD: C-terminal domain.

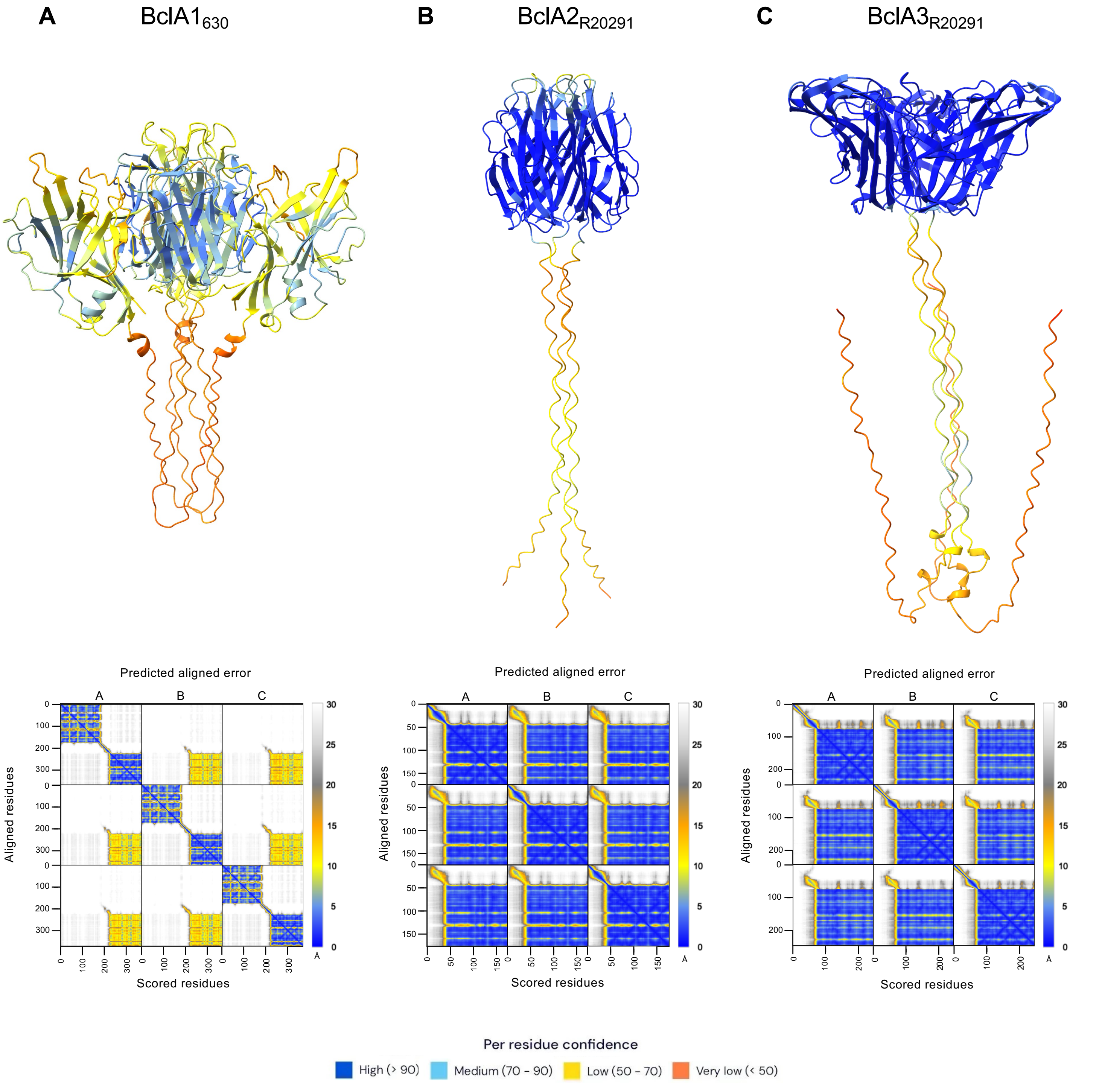


**Fig. S10. AlphaFold3 predicted structure of BclA_10GXY homotrimer.** Predicted structure of *C. difficile* **(A)** BclA1$_{630}$, **(B)** BclA2$_{R20291}$ and **(C)** BclA3$_{R20291}$ homotrimer complexes. Each model has its respective Predicted aligned error (PAE) plot below.

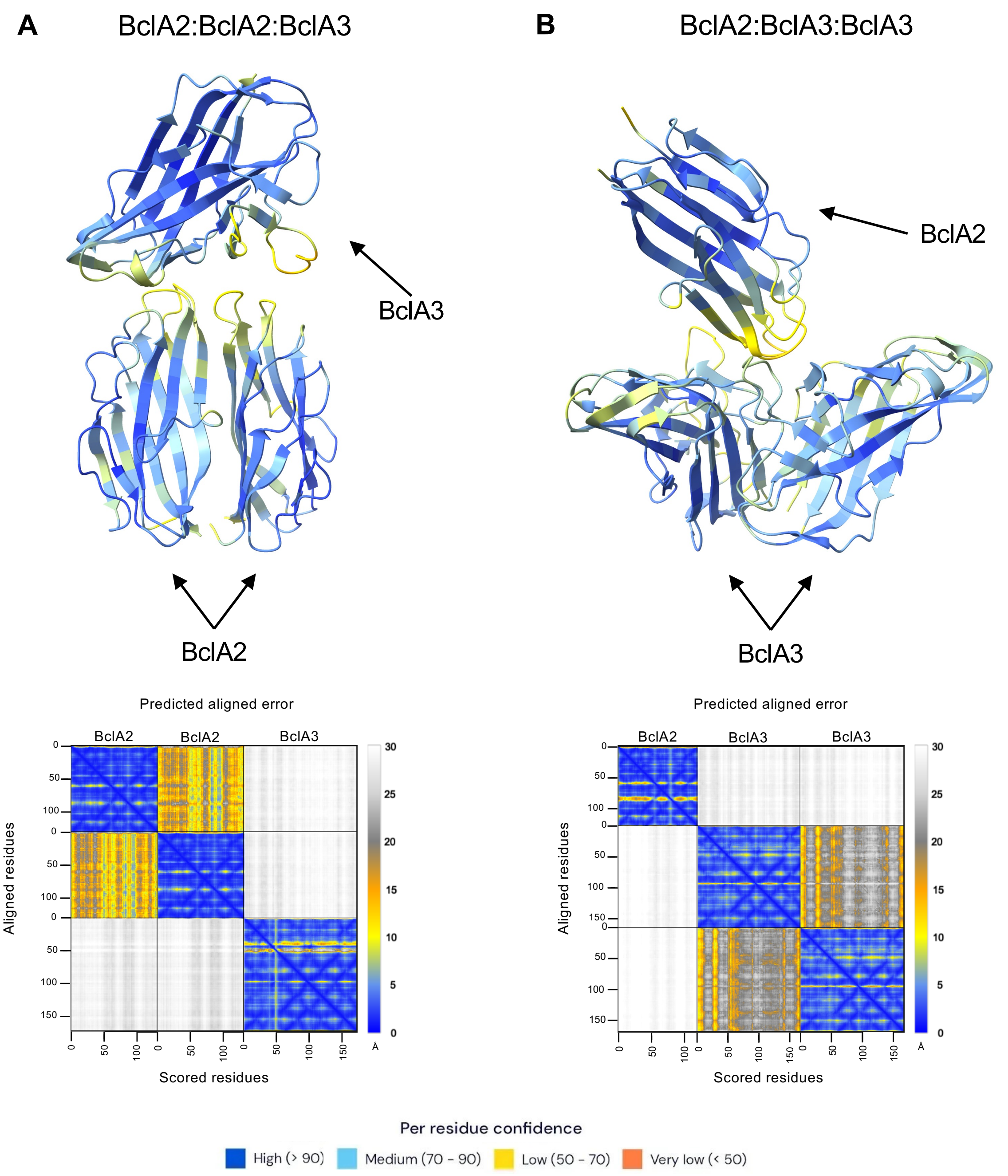


**Fig. S11. AlphaFold3 predicted CTD heterotrimer structures from R20291 strain.** Heterotrimer predicted model composed of **(A)** 2 units of BclA2_CTD and 1 unit of BclA3_CTD or **(B)** 1 unit of BclA2_CTD and 2 unit of BclA3_CTD. Each model has its respective Predicted aligned error (PAE) plot below.

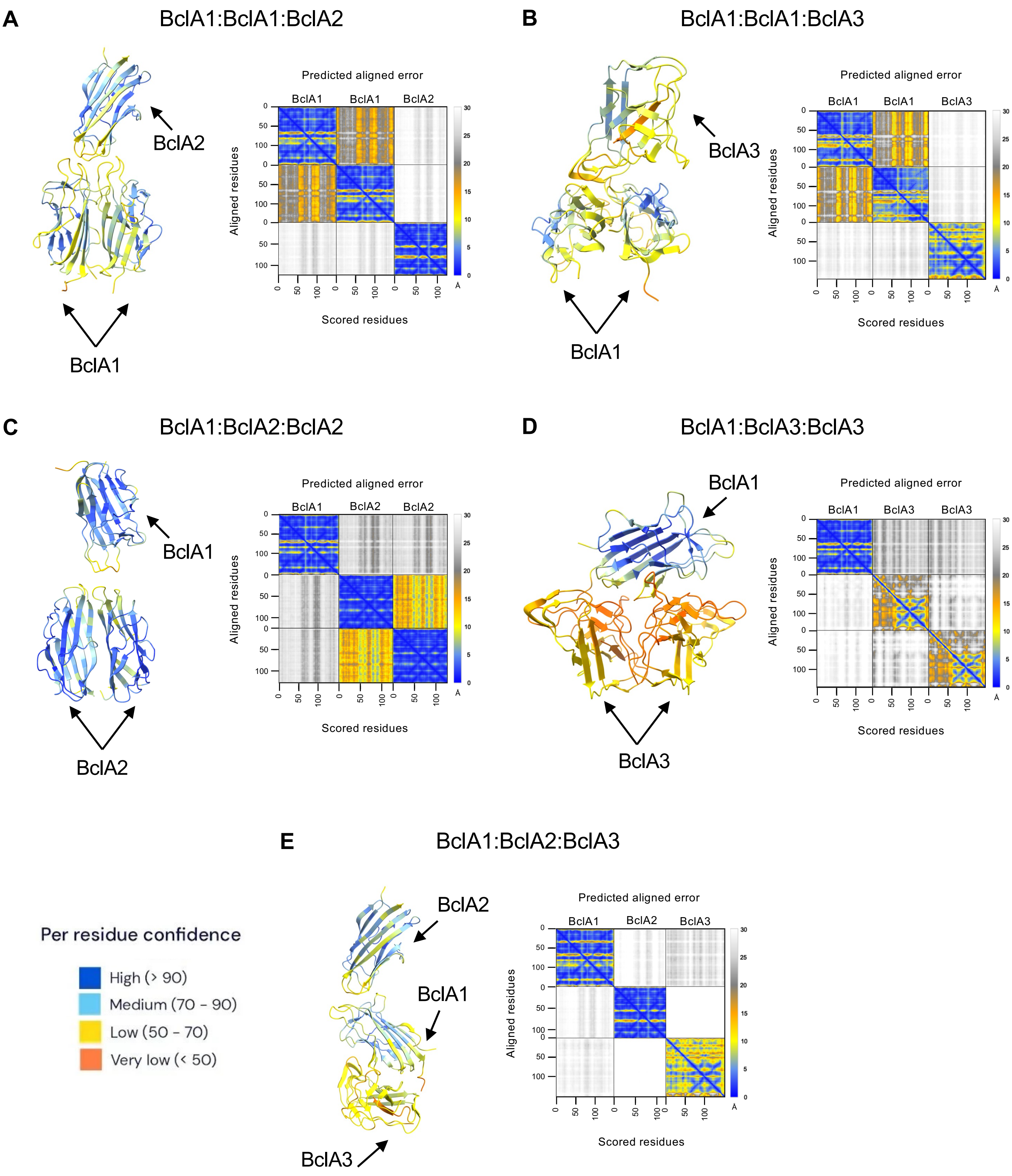


**Fig. S12. AlphaFold3 predicted CTD heterotrimer structures from 630 strain.** Heterotrimer prediction composed of **(A)** 2 units of BclA1_CTD and 1 unit of BclA2_CTD, **(B)** 2 units of BclA1_CTD and 1 unit of BclA3_CTD, **(C)** 1 unit of BclA1_CTD and 2 unit of BclA2_CTD, **(D)** 1 unit of BclA1_CTD and 2 unit of BclA3_CTD or **(E)** 1 unit of BclA1_CTD, 1 unit of BclA2_CTD and 1 unit of BclA3_CTD. Each model has its respective Predicted aligned error (PAE) plot.

**A**

```
BclA1              1 SQPVANFLVNAPSPQTLNNGDAITGWQTIIGNSSSITVDTNG--------     42
                                   ...|:|    |....:|.:..:..:|.|
BclA               1 --------------YAFNSG----GISLDLGINDPVPFNTVGSQFGTAIS     32

BclA1             43 -----TFTVQENGVYYISVSVALQPGSSSINQYSFAILFPILGGKDLAGL     87
                          ||.:.|.|.|  .::|.....::|:                |.||
BclA              33 QLDADTFVISETGFY--KITVIANTATASV----------------LGGL     64

BclA1             88 TTE------PGGGGVLSGYFAGFLFGGTTFTINNFSSTTV---GIRNGQS    128
                     |.:      ||.|..|....|..:....|......|...|   |:....:
BclA              65 TIQVNGVPVPGTGSSLISLGAPIVIQAITQITTTPSLVEVIVTGLGLSLA    114

BclA1            129 AGTAATLTIFRIADTVMT    146
                     .||:|::.|.::|
BclA             115 LGTSASIIIEKVA-----    127
```

**B**

```
BclA2              1 ANNAQFTVSSS----SLGNNTLVTFN---SSFINGTNITFPTSSTINLAV     43
                          :..:|.    .||.|..|.||   |.|  ||.|:...:.|..::.
BclA               1 -----YAFNSGGISLDLGINDPVPFNTVGSQF--GTAISQLDADTFVISE     43

BclA2             44 GGIYNVS-FGIRAILSLAGFMSITTNFNGVAQNN-----------FIAKA     81
                     .|.|.:: ....|..|:.|  .:|...|||....           .:.:|
BclA              44 TGFYKITVIANTATASVLG--GLTIQVNGVPVPGTGSSLISLGAPIVIQA     91

BclA2             82 VNTLTSSDVSVSLSFLVDARAAAVTLSFTFGSGTTGTSPAGYVSVYRIQ    130
                     :..:|::      ..||:.....:.||...|:..:       :.:.::.
BclA              92 ITQITTT------PSLVEVIVTGLGLSLALGTSAS-------IIIEKVA    127
```

**C**

```
BclA3              1 -----SQPVANFLVNAPSPQTLNNGDAITGWQTIIG---NSSSITVDTNG     42
                          |......||||    .|.|    ||  |::|   :...|.....|
BclB               1 IIPFASGTTPALLVNA----VLAN----TG--TLLGFGFSQPGIAPGVGG     40

BclA3             43 TFTVQENGVYYISVSVALQPGSSSINQYSFAILFPILGGKDLAGLTTEPG     92
                     |.|:              .||  .:..|:|      :..:|  |:.|.
BclB              41 TLTI--------------LPG--VVGDYAF------VAPRD--GIITS--     64

BclA3             93 GGGVLSGYFAGFLFGGTTFTINNFSSTTVGIRNGQSAGTAATLTIFRIAD    142
                         |:|:|:.        |......|.|.|:.......||:.|...:|.
BclB              65 ----LAGFFSA--------TAALAPLTPVQIQMQIFIAPAASNTFTPVAP    102

BclA3            143 TVMT----------------------------------------------    146
                     .::.
BclB             103 PLLLTPALPAIAIGTTATGIQAYNVPVVAGDKILVYVSLTGASPIAAVAG    152

BclA3            147 ---------    146

BclB             153 FVSAGLNIV    161
```

**D**

| Aligned sequence | Identity | Similarity |
|---|---|---|
| BclA1 - BclA | 31/168 (18.5%) | 51/168 (30.4%) |
| BclA2 - BclA | 29/149 (19.5%) | 53/149 (35.6%) |
| BclA3 - BclB | 33/209 (15.8%) | 50/209 (23.9%) |

**Fig. S13. Pairwise alignment (PSA) of *C. difficile* BclA_CTD with *B. anthracis* BclA/BclB_CTD.** EMBOSS Needle was used for PSA of **(A)** Cd_BclA1_CTD$_{630}$ with Ba_BclA_CTD **(B)** Cd_BclA2_CTD$_{R20291}$ with Ba_BclA_CTD and **(C)** Cd_BclA3_CTD$_{R20291}$ with Ba_BclB_CTD. **(D)** Percentage of identity of aligned sequences. CTD: C-terminal domain. *C. difficile* 630 genome used as reference was GenBank sequence: AM180355.1. *C. difficile* R20291 genome used as reference was GenBank sequence: CP029423. *B. anthracis* genome used as reference was GenBank sequence: AE016879.1.

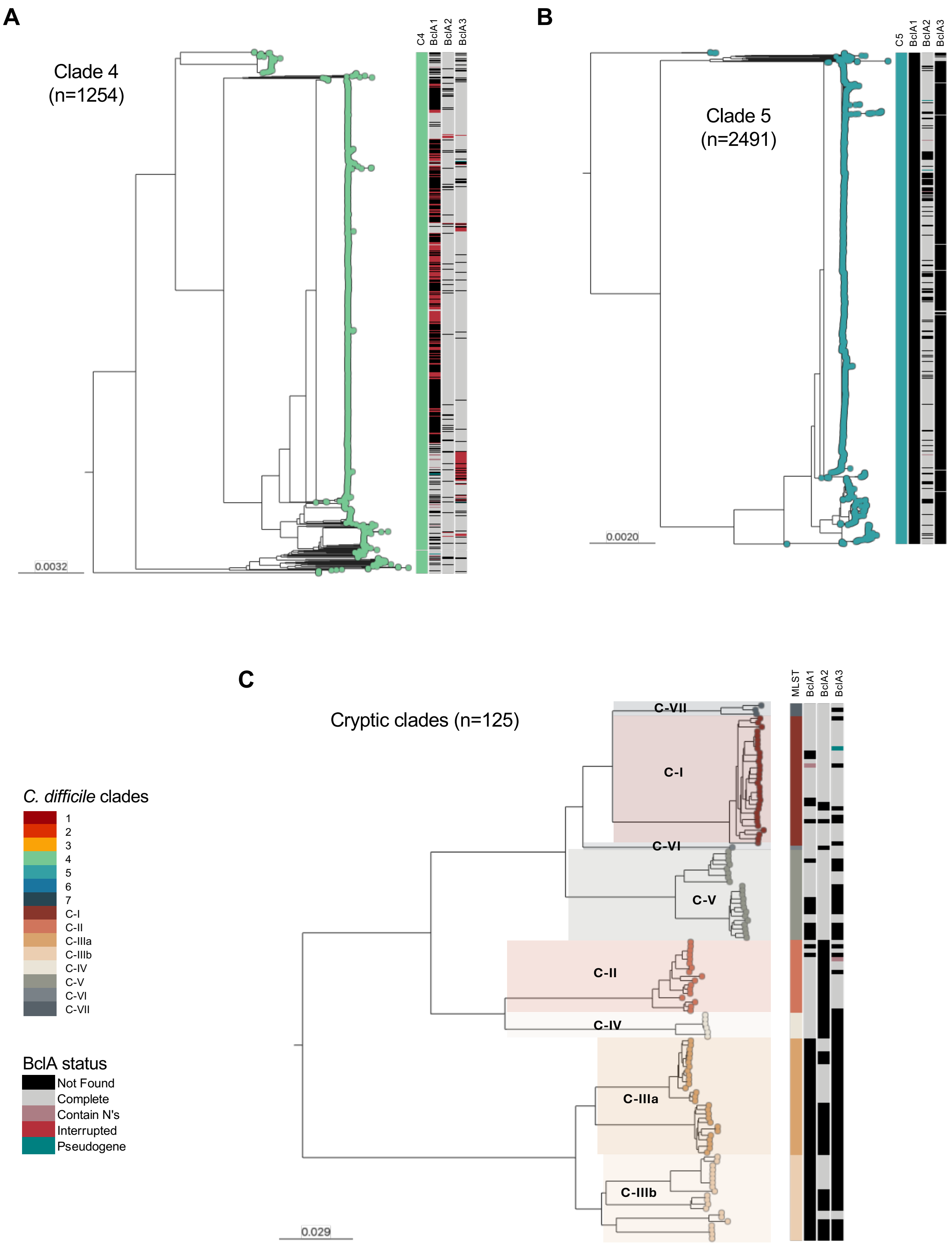


**Fig. S14. Distribution of BclAs in known diversity of *C. difficile*. (A)** Zoomed-in view of the Clade 4 subtree. **(B)** Zoomed-in view of the Clade 5 subtree. **(C)** Zoomed-in view of the cryptic clades subtree. Horizontal bars represent the state of BclA in a particular genome (not found, complete, containing Ns, interrupted, or pseudogenized).

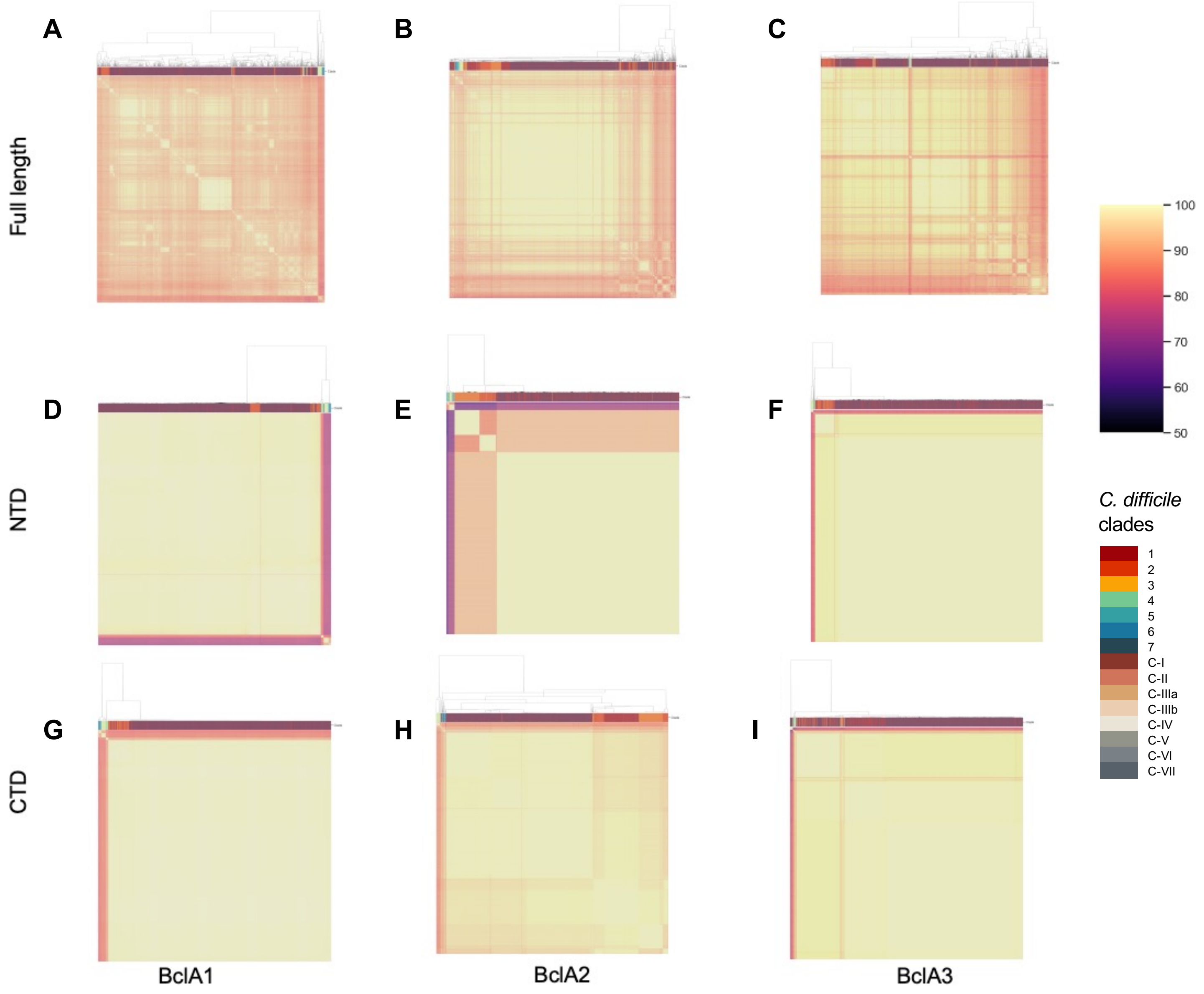


**Fig. S15. Average amino acid identity (AAI) analysis of BclA unique alleles.** Heatmap of the percentage of identity between all unique alleles of **(A)** BclA1, **(B)** BclA2 and **(C)** BclA3. AAI was performed for full length protein, NTD and CTD regions. N-terminal domain (NTD) and C-terminal domain (CTD).
